# A multifunctional type IA DNA/RNA topoisomerase with RNA hydrolysis and rRNA processing activities from *Mycobacterium smegmatis* and *Mycobacterium tuberculosis*

**DOI:** 10.1101/2020.07.13.199935

**Authors:** Phoolwanti Rani, Shashwath Malli Kalladi, Harsh Bansia, Sandhya Rao, Rajiv Kumar Jha, Paras Jain, Tisha Bhaduri, Valakunja Nagaraja

## Abstract

Topoisomerases maintain topological homeostasis of bacterial chromosomes by catalysing changes in DNA linking number. The resolution of RNA entanglements occurring in the cell would also require catalytic action of topoisomerases. We describe RNA topoisomerase and hydrolysis activities in DNA topoisomerase I (topo I) from mycobacteria. The interaction of topo I with mRNA, tRNA and rRNA suggested its role in some aspect of RNA metabolism; the enzyme participates in rRNA maturation via its RNA hydrolysis activity. Accumulation of rRNA precursors in a topo I knockdown strain and the rescue of rRNA processing deficiency in RNaseE knockdown cells by topo I expression, indicated the enzyme’s back-up support to RNases involved in rRNA processing. We demonstrate that the active site tyrosine of the enzyme mediates catalytic reactions with both DNA/RNA substrates, and RNA topoisomerase activity can follow two reaction paths in contrast to its DNA topoisomerase activity. Mutation in the canonical proton relay pathway impacts DNA topoisomerase activity while retaining activity on RNA substrates. The mycobacterial topo I thus exemplifies the resourcefulness and parsimony of biological catalysis in harnessing the limited chemical repertoire at its disposal to find common solutions to mechanistically-related challenges of phosphodiester breakage/exchange reactions in DNA and RNA that are essential for cell survival.

## Introduction

Topoisomerases, the “Magicians of the DNA world” [1], are enzymes dedicated to resolving topological hurdles encountered during copying, processing and exchange of biological information, *viz*, replication, transcription and recombination. In this role, topoisomerases catalyse supercoiling/relaxation, catenation/decatenation and knotting/unknotting reactions in DNA. All topoisomerase reactions share the following steps: introduction single-strand nicks (Type I) or double-strand breaks (Type II) in DNA, strand passage or strand rotation, and resealing of the broken DNA ends [1-6]. The occurrence, number and distribution of different classes of DNA topoisomerase vary among organisms depending on cellular context and/or complexity [1, 4]. While the roles of DNA topoisomerases in modulating DNA topology have been extensively studied, by contrast, topological problems associated with RNA transactions have been underappreciated, presumably because most RNA is single-stranded. Thus, topoisomerase reactions in RNA substrates have received less attention than analogous reactions in their DNA counterparts. The demonstration of an RNA topoisomerase activity in *E. coli* topoisomerase III (topo III) set the stage for exploring dual activities in other systems [7]. The activity on RNA was subsequently reported with vaccinia virus topoisomerase and human topoisomerase I, which are type IB enzymes unlike the type IA *E. coli* topo III [8]. RNA topoisomerase activity was also documented in human top 3β and in organisms from all three domains of life [9]. Nevertheless, a variety of topological changes in RNA substrates that these enzymes can bring about, the mechanistic paths of these reactions, and their potential role in other aspects of RNA metabolism are largely unknown.

We have addressed some of these questions by examining the RNA topoisomerase activity of topo I, the sole type IA enzyme in *Mycobacterium tuberculosis* and *M. smegmatis* [10-12]. Our earlier studies indicated that the enzyme in both species is essential for cell survival [13, 14]. This is not surprising, as there are no other type IA enzymes in these bacteria to shoulder the responsibilities [12]. Notably, in addition to being a DNA relaxase, topo I appears to participate along with DNA gyrase at the Ter site to carry out decatenation and chromosome segregation [11, 15-17]. Here we describe a new activity other than topoisomerisation for the topo I from mycobacteria. In addition to RNA topoisomerase activity, the enzyme participates in processing of precursor RNAs generated from rRNA by its RNA hydrolysis activity. The catalytic versatility of topo I permits mycobacteria to perform multiple mechanistically-related but functionally distinct phosphoryl transfer reactions in DNA and RNA using a single protein.

## Result

### RNA binding, cleavage, and formation of covalent intermediates by TopoI

In our experiments while characterizing the DNA relaxation activity of TopoI, we observed inhibition by addition of tRNA (Supplementary Figure S1A). The apparent competition by tRNA was consistent with the ability of *M. tuberculosis* and *M. smegmatis* topo I (henceforth MtTopoI and MsTopoI respectively) to bind and cleave tRNA and single stranded (ss) DNA (Figure 1A and B; Supplementary Figure S1B and C). The cleavage activity on tRNA was competed by 32-mer single stranded DNA (Supplementary Figure S1D), indicating that both tRNA and ssDNA are potential substrates for the enzyme. Being a type IA topoisomerase, the enzyme forms 5’ phospho-tyrosine adducts with DNA as the cleavage intermediate during relaxation [11]. Formation of the 5’phospho-tyrosine intermediates was also evident with the tRNA substrate (Figure 1C). Furthermore, the enzyme bound rRNA and mRNA (Supplementary Figure S1 E and F), suggesting that a broad range of RNAs are likely to be potential substrates for topo I. Interaction of the enzyme with intracellular RNA was revealed by RNA immunoprecipitation (RIP) (Figure 1D; Supplementary Figure S2 A and B); its association with rRNA, a few of the mRNA, and a tRNA locus was verified by RIP-qPCR (Figure 1D). Together, these data suggest the potential participation of the enzyme in one or more RNA transaction events.

**Figure 1.**
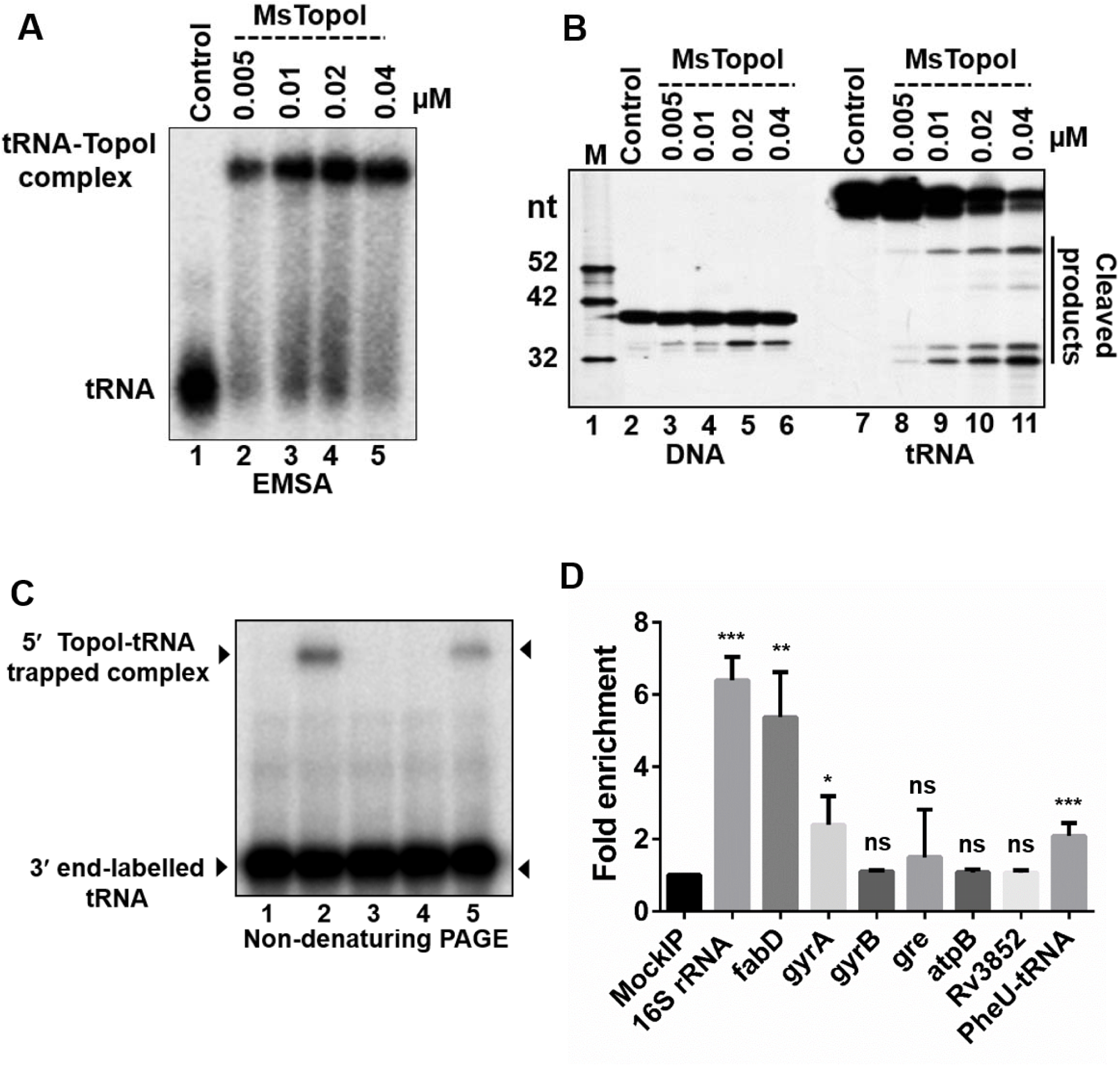
Topo I interaction with RNA *in vitro* and *in vivo*. (A) EMSA with itRNA. Lane 1, 5’-end labeled initiator tRNA (itRNA), lanes 2-5, itRNA incubated with indicated concentrations of MsTopoI and the products were analysed on non-denaturing PAGE. (B) Cleavage assays. 5’-end labelled ss DNA and itRNA were incubated with indicated concentrations of MsTopoI and the products were visualized on 8 M urea-6% PAGE. M, oligonucleotide markers; lanes 2 and 7, ssDNA/itRNA controls. (C) Covalent complex of topo I-itRNA. The covalent complexes were generated by arresting the reactions (with 0.5% SDS) of 3’-end labelled itRNA with the enzyme and separating the products on non-denaturing 6% PAGE. Lane 1, itRNA alone. Lane 2, enzyme-itRNA complex. Lane 3, complexes treated proteinase K. Lane 4 and 5, competition with 5 pmol of specific and non-specific ss DNA, respectively (Materials and Methods). (D) RNA immunoprecipitation (RIP)-qPCR. Complexes of RNA with topo I were immunoprecipitated and processed as described in Materials and Methods and Supplementary Figure 2A. The data represented is mean ± SEM of three independent samples. P values were determined by unpaired t test using GraphPad Prism 6.0. P value≤0.05 was considered as significant. ns-non-significant, * <0.05, ** <0.01, and *** <0.001.

### RNA processing function

The interaction of topo I with various types of RNA seen in Figure1 and Supplementary Figure 1, led us to consider enzyme’s participation in the processing of precursor RNAs for the following reasons. 1. Binding, cleavage, and covalent complex formation with RNA by the enzyme are unlikely to be artefactual. 2. The rRNA processing enzymes are underrepresented in mycobacteria (Supplementary Table S1), 3. The scarcity in processing RNases in the organism may warrant the contribution by a back-up function under certain conditions. Transcription of the rRNA operon and several tRNA operons driven by their respective promoters generates polycistronic precursor RNAs, which are then processed to yield mature RNAs. Total RNA isolated from the cells contains, in addition to mature RNAs, polycistronic RNA as well as partially processed precursor 16S, 23S and 5S rRNAs. This population of RNA molecules was treated with MsTopoI, and Northern hybridization analysis was carried out with the probes depicted in Figure 2A. A decrease in 5’ and 3’precursor region of both 16S and 23S rRNA was observed, suggesting that they are substrates for processing by topo I (Figure 2A). When the *in vitro* transcribed 16S rRNA containing the precursor region was treated with wild-type topo I or the active-site mutant Y339F [18] (Figure 2B), cleaved products were seen only with wild-type topo I. Similarly, when the 16S and 23S intergenic precursor region was amplified, *in vitro* transcribed, and treated with MsTopoI, cleaved products were observed in the case of the wild-type enzyme but not with the active-site mutant form (Figure 2C). These data indicate that the catalytic tyrosine residue is required for the hydrolysis of RNA. To examine tRNA-hydrolysing activity by the enzyme, one of the operonic tRNA substrate was used (Materials and Methods). As with the ribosomal RNA precursor, cleavage of the tRNA precursor was seen with MsTopoI but not with the Y339F mutant (Figure 2D). Similar results were obtained when 16S rRNA with precursor region, intergenic region between 16S and 23S rRNA and operonic tRNA were treated with MtTopoI (Figure 2E).

**Figure 2.**
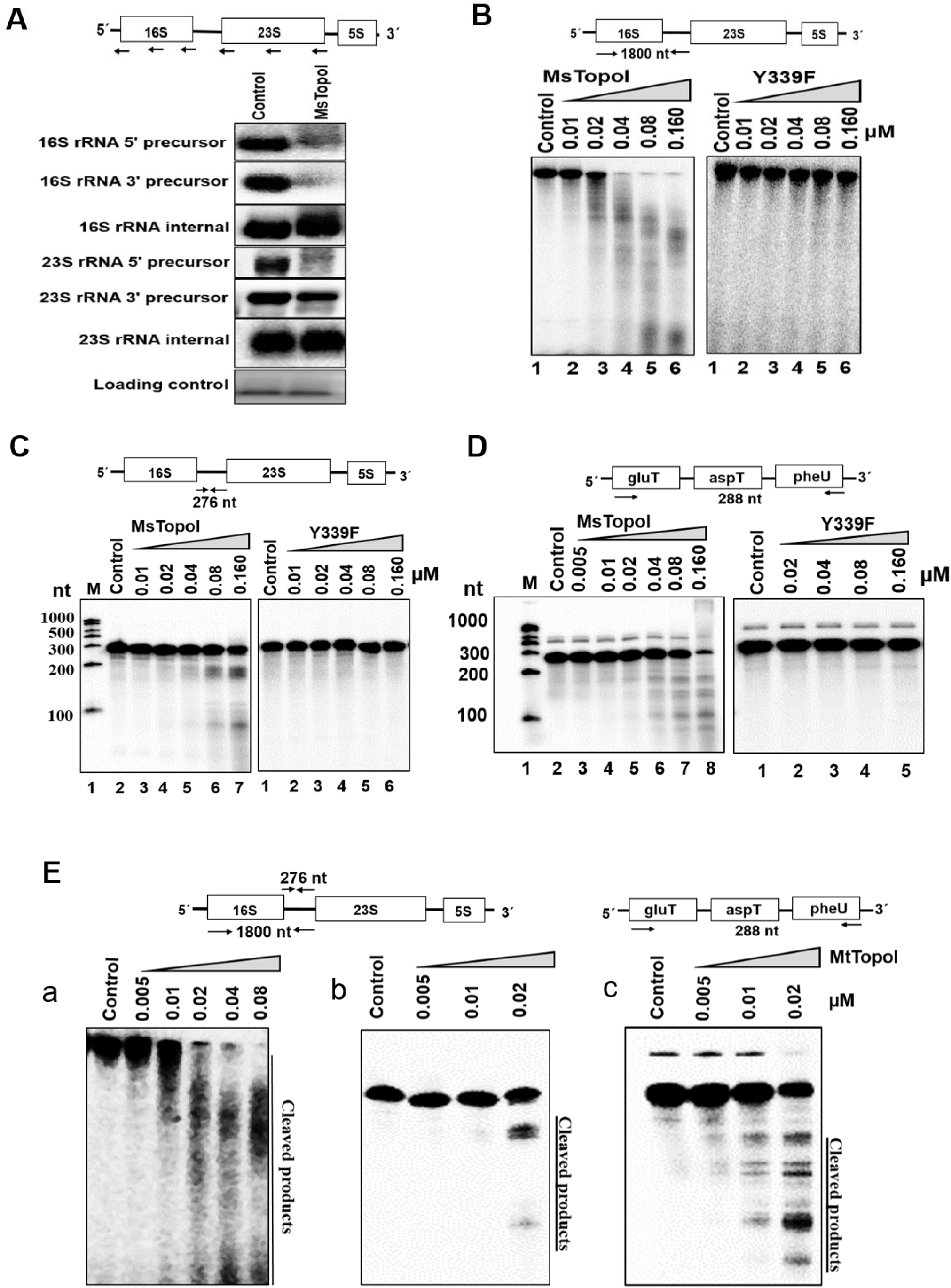
RNA processing assays. (A) MsTopoI treated RNA (isolated from ΔRNaseR strain of *E. coli*) probed for precursor and internal regions of 16S and 23S rRNA as shown in the Figure and described in Materials and Methods. The loading control is from the same gel stained with EtBr before transferring to the membrane for Northern analysis. Schematic representing the probes for internal and precursor regions of 16S and 23S rRNA are shown at the top. (B) *In vitro* transcribed 16S rRNA with precursor region (1800 nucleotides, schematic shown above) treated with MsTopoI and Y339F and the products analysed by 8 M-18% PAGE. (C) *In vitro* transcribed intergenic region between 16S and 23S rRNA was treated with indicated concentrations of MsTopoI and Y339F; the products were analysed by 8 M urea-6% PAGE. (D) *In vitro* transcribed tRNA (gluT-aspT-pheU) was treated with MsTopoI and Y339F and analysed by 8M urea-6% PAGE. (E) RNA processing with MtTopoI. *In vitro* transcribed 16S rRNA with precursor region (1800 nt, a), 16S-23S intergenic region (276 nt, b), and tRNA (288 nt, c) were digested with MtTopoI and analysed as in B, C, D respectively.

To evaluate the involvement of topo I in rRNA maturation *in vivo*, analysis was carried out in topo I knockdown cells. For this, a topo I conditional knockdown strain was generated using the CRISPR interference system (Materials and Methods, [19]) and repression of topo I expression was confirmed by immunoblots (Figure 3A). If topo I participates in RNA processing *in vivo*, there should be an accumulation of precursor rRNA in topo I-depleted cells. Accumulation of 16S precursor rRNA in the topo I depleted strain was detected by RT-qPCR and confirmed by Northern blots (Figure 3 B-D, Supplementary Figure 3A and B).

**Figure 3.**
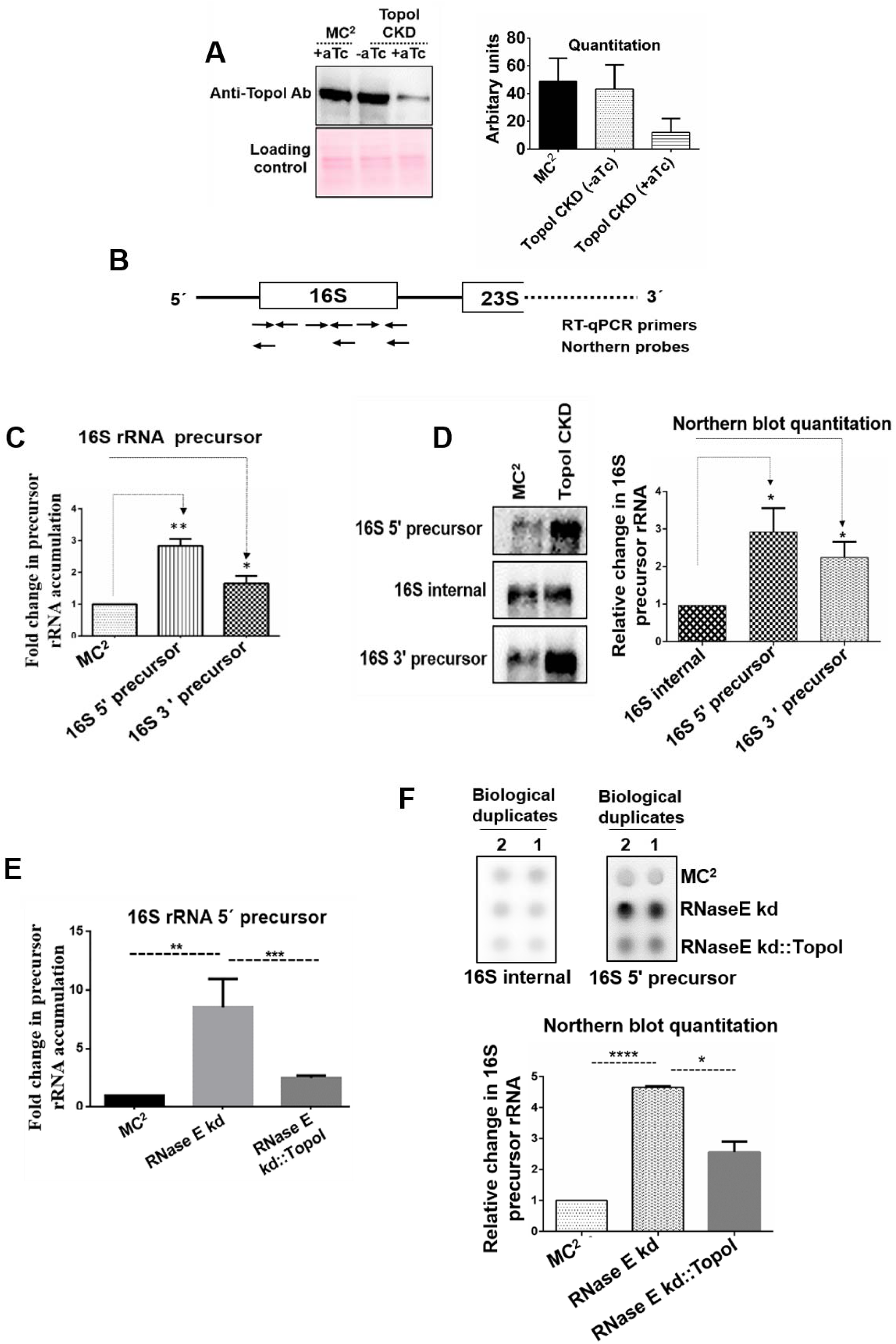
rRNA processing by topo I *in vivo*. (A) Immunoblot analysis showing reduction in topo I levels in topo I CKD strain. MC^2^, *M. smegmatis* strain depicting normal topo I levels. The same blot stained with ponceau is shown as a loading control. Right panel shows quantitation of blot (using Image J) of three independent experiments. (B) Schematic depicting primers and probes (Supplementary Table S2) to assess precursor rRNA accumulation. (C) RT-qPCR. The data from real-time PCR performed with RT primers for precursor and internal regions represented after normalizing with the internal region. Error bar represents mean ± SEM of three independent experiments. Paired t tests were used to determine P values. *<0.05 and ** <0.01. (D) Northern analysis showing precursor rRNA accumulation in topo I CKD strain compared to *M. smegmatis* MC^2^. Quantitation is shown on the right panel. (E and F) Complementation of rRNA processing defect by TopoI. RT-qPCR (panel E) and Northern dot blots (panel F) analysis of RNA isolated from RNaseE knockdown (RNaseE kd) and topo I complemented (RNaseE kd::TopoI) strains as described in Materials and Methods. The quantitation is shown in the lower panel of Figure F. To calculate relative fold changes, precursor 16S rRNA band (5’ or 3’) pixel values were normalised by the internal 16S rRNA band values and subsequently expressed relative to MC^2^. The error bar in RT-qPCR data represents the mean ± SEM of three independent experiments. P values in D, E and F were determined by unpaired t test using GraphPad Prism 6.0. *<0.05, ** <0.01 and *** <0.001.

### Functional cooperation between ribonucleases and topo I in RNA processing

Only 7 RNases are found in mycobacteria that participate in precursor rRNA processing in contrast to the larger number (11) found in *E. coli* (Supplementary Table S1). Given this relative paucity in rRNA maturases, the rRNA hydrolysing activity of topo I seen above appeared to be significant, and the potential role of MsTopoI as a *de facto* RNA processing enzyme in mycobacteria becomes functionally relevant.

Hence, to further examine whether MsTopoI is involved in rRNA processing, an RNaseE knockdown strain (RNAseE kd) was generated in which its expression is downregulated by promoter replacement and control by two-repressor system as described (Materials and Methods, [20]). RNaseE is an essential endoribonuclease required for maturation of precursor region of 16S, 23S, and 5S rRNA in *M. smegmatis* [21]. Reduction in RNaseE levels in the conditional knockdown strain was determined by RT-qPCR (Supplementary Figure S4A), and the resultant accumulation of precursors of 16S and 23S rRNA was assessed by RT-qPCR and Northern dot-blots (Supplementary Figure S4 B). With about 30% reduction in RNaseE (Supplementary Figure S4A), 5-15-fold increase in 16S 3’, 16S 5’ and 23S 5’ precursor accumulation was observed respectively (Supplementary Figure S4B and C). When topo I was expressed in the RNaseE knockdown strain (Materials and Methods; Supplementary Figure S4D), partial rescue of the rRNA maturation defect was seen (Figure 3E and F). A substantial decrease in precursor accumulation (∼8.5 fold to ∼3 fold compared to *M. smegmatis* MC^2^, Figure 3E) upon near restoration in topo I levels in RNaseE knockdown background points to its participation in processing (Figure 3E and F). Thus, the complementation of RNaseE deficiency by topo I suggests a supernumerary role for topo I as an RNA maturase in mycobacterium, and perhaps other microorganisms with a deficit of RNases devoted to this task.

### RNA topoisomerase activity of mycobacterial topo I

RNA topoisomerase activity was not detected with *M. smegmatis* and *M. tuberculosis* topo I in a previous study [9]. Although RNA topoisomerase activities have been described with type IA topoisomerases, lack of such activity with the sole type IA enzyme in mycobacteria was rather surprising. Since, we could detect binding and cleavage of RNA (Figure 1), and also RNA processing function (Figure 2 and 3), we re-evaluated the RNA topoisomerase activity in mycobacteria. RNA topoisomerase activity of DNA topoisomerases was inferred in earlier studies by demonstrating their ability to tie knots in circular RNA molecules [7, 9, 22]. Hence, a circular RNA substrate (Figure 4A) containing two complementary regions separated by a spacer sequence (Materials and Methods; [7, 22]) was incubated with MsTopoI, MsTopoI Y339F, and MtTopoI. The conversion of RNA circle to RNA knot(s) was observed with the MsTopoI and MtTopoI but not with the mutant (Figure 4B and C, Supplementary Figure 5A). The difference between wild type and active-site mutant is consistent with the expected knotting pathway, namely, phosphodiester cleavage, RNA strand passage and religation. Next, we examined the ability of MsTopoI to simplify RNA topology. An entangled RNA substrate was prepared as described in Materials and Methods, and subjected to topo I reaction. Products with higher electrophoretic mobility than the substrate were formed (Supplementary Figure 5B), as expected for the removal of RNA crossings by the enzyme.

**Figure 4.**
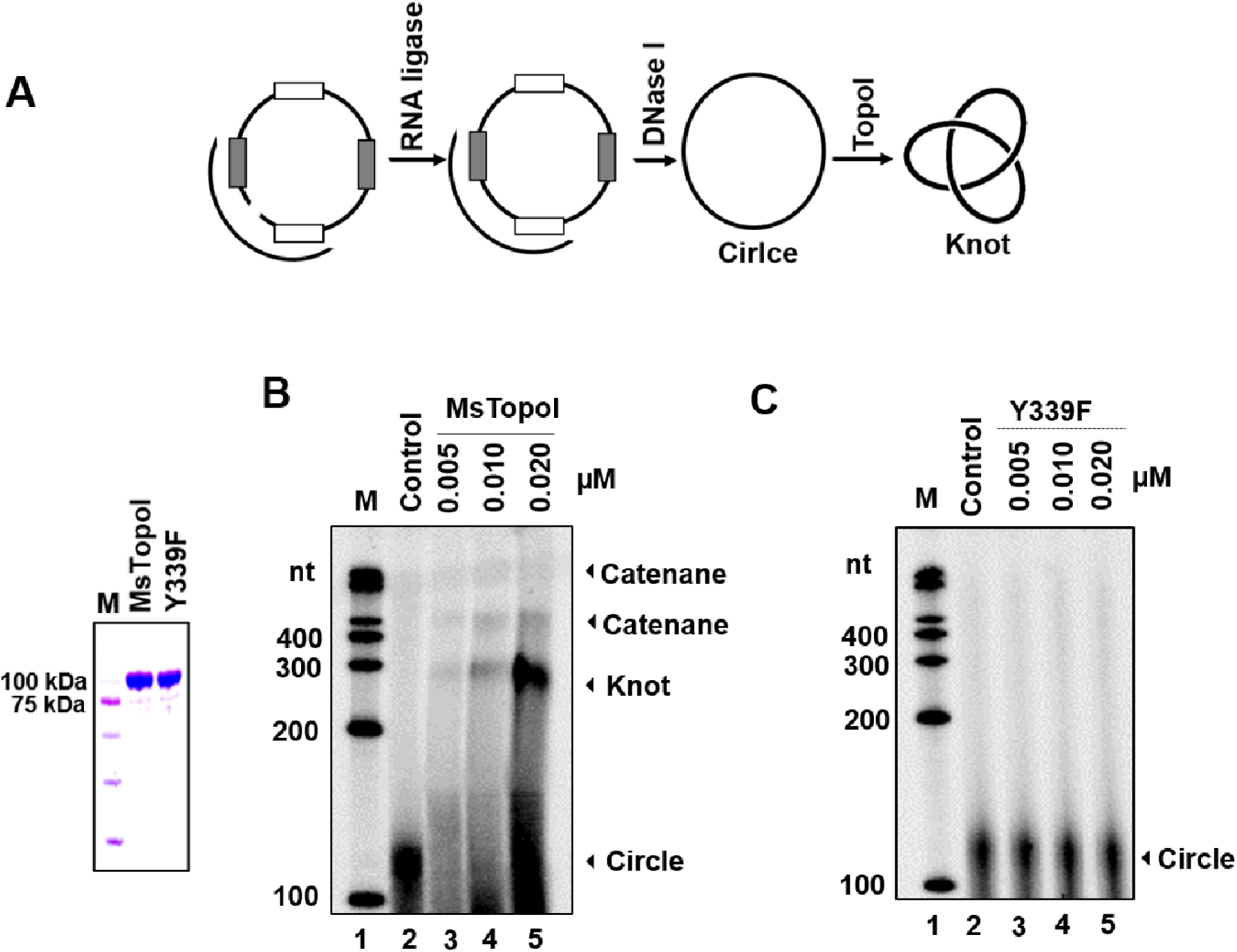
RNA knotting. (A) Schematic of the knotting experiment. The design of the substrate is as described [7, 22], Materials and Methods). The empty and filled rectangles on the RNA depict the complementary regions (14 nucleotides) separated by 12-nucleotides spacer sequences (Supplementary Table S2). (B and C) RNA knotting. The circular RNA substrates were incubated with MsTopoI (B) and Y339F mutant (C) and the products were analysed by 8 M urea-15% PAGE. The assignment of circle/knot/catenane are based uppon the previous studies [7, 22]. Left panel is a Coomassie-stained gel showing MsTopoI and Y339F mutant.

In all the studies on RNA topoisomerases so far, not all the individual steps of the topoisomerase reaction have been investigated and the conversion of the circular RNA into knots was the only assay employed. For an enzyme to be an RNA topoisomerase, it must bind and cleave RNA, pass an intact strand through the opening, and ligate the broken ends. While the results shown in Figure 1 and Supplementary Figure S1 establish the first two steps of the reaction, for probing RNA strand passage, we adapted a DNA strand-passage assay that we developed earlier [23]. The central cavity of the enzyme can accommodate either a single-stranded or double-stranded nucleic acid [24-29]. During DNA relaxation, the unbroken strand (T segment) can enter the central cavity or exit from it through the cleaved DNA gate [24-26, 30]. In the original assay for DNA strand passage, pUC18 DNA was incubated with the topo I to occupy the enzyme cavity. The gate remains closed by the intact site-specific oligo (G segment) bound to topo I before oligonucleotide cleavage. The gate is opened by oligonucleotide cleavage by the enzyme [23]. Anti-topo I clamp-closing monoclonal antibodies (1E4F5) were used to lock the enzyme gate, trapping the plasmid in the enzyme cavity [31], either before or after cleavage of the oligonucleotide. The extent of escape of T-segment DNA from the central cavity of the enzyme provides a measure of strand-passage activity [23]. With labelled tRNA as the reporter (Materials and Methods; Figure 5A), the reduction in retained radioactivity following topo I addition, attests to its RNA strand-passage competence (Figure 5B). The carboxyl-terminal domain (CTD) of topo I harbours the DNA strand passage activity [16, 23]. Three stretches of basic residues within the CTD make non-equivalent contributions towards full activity [23]. RNA strand-passage activity was abolished by deletion of the CTD (Figure 5B), indicating its indispensable role for RNA topoisomerase activity as well. Furthermore, deletion of only the third basic stretch reduced strand passage by about four-fold while deletion of both the second and third stretches led to complete loss of activity (Figure 5B). Deletion of the first basic stretch, by contrast, resulted in only a partial loss of activity.

**Figure 5.**
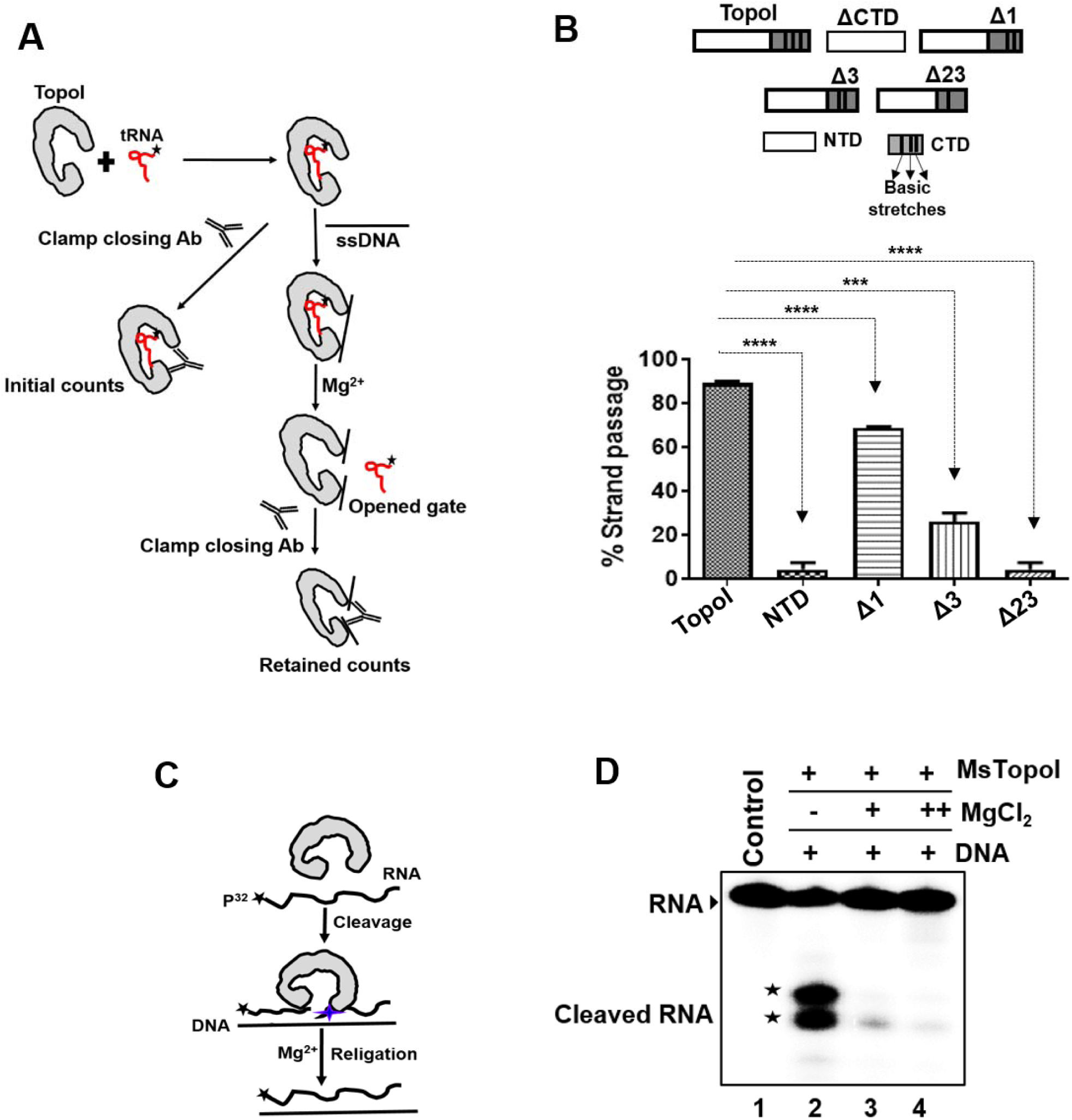
Strand passage and religation. (A) Schematic of strand passage experiment. tRNA captured in the cavity of the enzyme and enzyme is locked by clamp closing antibodies before and after strand passage as described in Materials and Methods and Results section. (B) Strand transfer reaction with MsTopoI and its variants. The CTD is depicted in grey; the three basic stretches and their deletions are as shown. The extent of strand passage was determined by measuring the initial and final radioactivity retained on the filters as depicted in A and as described in Material and Methods. The error bars represent standard error mean across the three measurements. P values were determined as in Figure 1. (C) Schematic of RNA religation. Cleaved RNA fragments were annealed to complementary DNA and religation initiated by Mg^2+^. (D) Religation assay. Lane 1, labelled RNA; lane 2, Cleaved RNA after incubation with MsTopoI; lanes 3 and 4, joining of cleaved RNA strands in the presence of 5 mM and 10 mM Mg^2+^respectively.

Next, RNA religation activity (second transesterification reaction following strand passage) of the enzyme was assayed using a synthetic substrate in which the two ends of a linear RNA were brought into proximity using a short complementary DNA as an anchor (Materials and Methods; Figure 5C). The cleaved fragments seen in the reaction without Mg^2+^ were religated by topo I in a metal ion (Mg^2+^)-dependent manner (Figure 5D), indicating a Mg^2+^ dependency for the second transesterification reaction, as observed with DNA topoisomerase reactions.

### DNA topoisomerase *vs* RNA topoisomerase and RNA topoisomerase *vs* hydrolysis: mechanistic considerations

Irrespective of whether a DNA or RNA substrate is used, nucleophilic attack on the scissile phosphodiester bond is the first catalytic step for topoisomerase activity that is followed by strand passage and religation of the broken ends of the DNA or RNA. Thus, during DNA or RNA cleavage by mycobacterial topoisomerase, and concomitant formation of the covalent adduct, Y339 (of MsTopoI) and the 3’-hydroxyl group of the scissile phosphate would serve as the nucleophile and leaving group. A nearby water molecule would deprotonate the nucleophile, thereby activating it for the attack on the scissile phosphodiester bond [32]. Protonation of the leaving group results in the formation topo I-DNA or RNA phosphotyrosine intermediate and cleaved DNA or RNA fragments. The reaction is thought to proceed via acid-base catalysis assisted by a His-Asp-Glu catalytic triad for DNA topoisomerase reaction (Figure 6A; reaction path 1) [32, 33]. The presence of the 2’-hydroxyl group in RNA provides for an additional alternative cleavage mechanism (Figure 6A; reaction path 2). Analogously, the religation reaction, which is chemically a reversal of cleavage, has two mechanistic options for the RNA substrate but only one for the DNA substrate (Figure 6B). A disruption in the proton relay at a critical point would arrest the DNA topoisomerase reaction but should allow RNA topoisomerase activity via reaction (2). Hence, we examined whether disrupting the normal proton relay within the catalytic triad has differential impact on the cleavage/religation activities in DNA versus RNA substrates.

**Figure 6.**
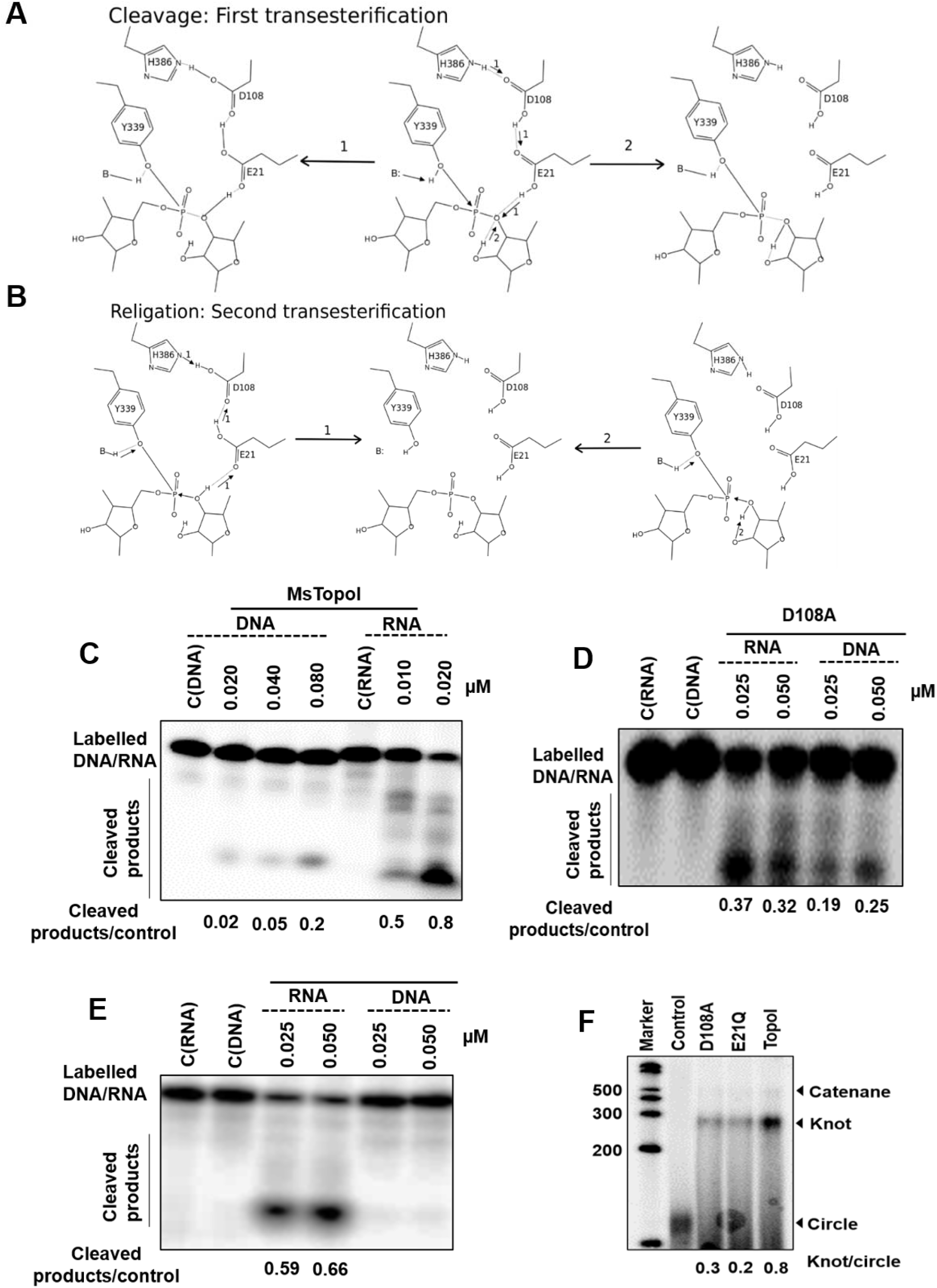
Mechanism of topo I-mediated reactions. (A) A nearby water molecule deprotonates the nucleophile (OH of Tyrosine-339 in MsTopoI) activating it for the attack on the scissile phosphodiester bond. Protonation of the leaving group (O of RNA 5’ to the site of cleavage) results in the formation of a topo I-RNA phosphotyrosine intermediate and a cleaved RNA fragment. The proximity of the 2’-OH to the leaving group in RNA would also allow pathway 2 for protonation of the leaving group. In pathway 1, the E21 side chain provides a proton to the leaving group which in turn extracts a proton from the H386 side chain via a proton relay through D108. In pathway 2, the proximal 2’-OH group provides a proton to the leaving group directly. (B) Second transesterification reaction. Deprotonation of the nucleophile (3’-OH of RNA) activates it for the attack on the topo I-RNA phosphotyrosine bond. The proximity of the 2’-OH to the nucleophile in RNA would allow two pathways for deprotonation of the nucleophile, depicted by marked arrows. Pathway 1 is the reversal of proton relay depicted in A. In pathway 2, the proximal 2’-oxygen extracts a proton from the nucleophile directly. Protonation of the leaving group (side-chain O of Tyrosine-339) by a nearby water molecule results in the religation of RNA and the release of TopoI. (C-E) First transesterification (cleavage) reaction. 5’-end labelled 28-mer DNA and RNA treated with MsTopoI (C), D108A (D) and E21Q (E) respectively. The products formed were analysed by 8 M urea-12% PAGE. (E) Assays with the circular RNA substrate (Materials and Methods, and Figure 4). The products were visualized by 8 M-15% PAGE. C, no enzyme.

Prior studies with MtTopoI and *E. coli* topo I showed that mutating the His-Asp residues of the triad (His-Asp-Glu) do not abolish the DNA cleavage of these enzymes [32, 33]. Presumably a suitably positioned water molecule is able to rescue the proton relay. However, the terminal Glu residue of the triad is critical for both the cleavage and religation steps. Substitution of this residue by Ala or Gln in MtTopoI results in undetectable DNA cleavage or DNA relaxation activity [32]. The results of the effect of the mutations in the catalytic triad in MsTopoI are presented in Figure 6C-F, and Supplementary Figure S6. MsTopoI and MtTopoI enzymes have a high degree of similarity; the active-site and catalytic-triad residues are conserved (Supplementary Figure S7 and S8). The substitution in the second amino acid of the catalytic triad (D108A) did not affect DNA cleavage by MsTopoI, as anticipated from published results (Figure 6C and D, [34]). However, the mutant had no detectable DNA relaxation activity because of the loss of religation activity (Supplementary Figure S6A and B; [34]). Strikingly, the E21Q variant of MsTopoI (terminal residue of proton relay), which has no DNA cleavage/religation activity, was active in the RNA cleavage and knotting reactions (Figure 6C, E and F, and Supplementary Figure S6A and C). While the cleavage activity was strong -(∼66% conversion of the input substrate to cleaved product; Figure 6E), the knotting activity was ∼4-fold lower than that of the wild-type MsTopoI (Figure 6F). Thus the 2’-hydroxyl group is able to rescue the absence of D108 and E21 of MsTopoI, efficiently in RNA cleavage and weakly in RNA religation.

The mechanism shown in Figure 6A is also applicable to RNA hydrolysis by topo I during RNA processing. The active-site tyrosine variant of MsTopoI (Y339F) showed no detectable DNA and RNA hydrolysis (Figure 7A and B). Here cleavage is likely followed by hydrolytic release of the covalently bound enzyme (Figure 7C). Utilization of water as the nucleophile by the topoisomerase would be analogous to the RNaseH reaction [35]. In addition to the transesterification reactions that topoisomerases catalyse, human topoisomerase I and vaccinia topoisomerase I are also able to carry out *in vitro* hydrolysis of the phosphotyrosyl bond in the cleaved intermediate, albeit at a low rate [36, 37]. Hydrolysis of the 3’-phosphotyrosyl bond by the human topoisomerase I (type IB) is pH-dependent [36]. To investigate the effect of pH on the relative abundance of transesterification and hydrolysis products, MsTopoI reactions were performed with 3’-end labelled DNA and RNA substrates in the pH range of 6.0 to 9.0 (Materials and Methods). With RNA, from 6-7.5 pH, hydrolysis was strongly favoured over covalent complex formation (Figure 7D). That the reaction is catalysed by the enzyme is apparent as hydrolysis is seen at the lower pH; if it were a general degradation it would have increased at higher pH. In contrast, the DNA covalent complex was stable in the entire pH range tested (Figure 7E). These results suggest that pH is one of the conditions that could influence the balance between the alternative cleavage products when the topoisomerase acts on RNA substrates. Although intracellular pH is thought to be tightly regulated, now it is apparent that the pH can change by about two units in different mycobacteria [38]. *M. tuberculosis* has found several uncanny ways to survive inside the host. It adapts to the acidic environment of host niche to survive in the acidic phagolysosomes of activated macrophages [39] by switching over metabolic status [40, 41]. The enzymes such as acid phosphatases and transglutaminases have the ability to switch to different enzyme activities at different pH [42, 43]. Thus, we envision a scenario in which mycobacterial topo I switching from hydrolysis to topoisomerisation and vice versa concomitant with the transient changes in pH especially under conditions of stress.

**Figure 7.**
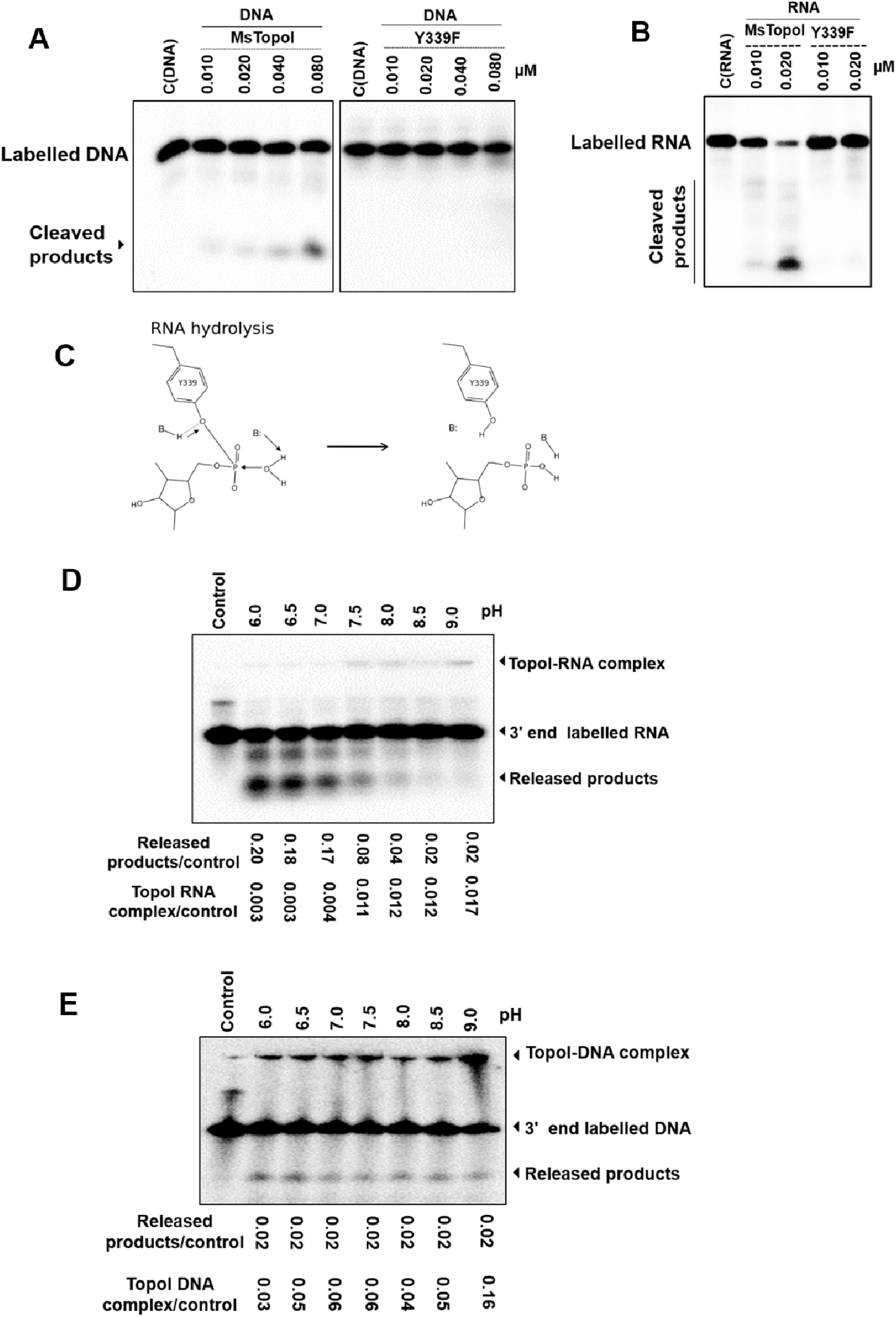
RNA hydrolysis activity of topo I. (A and B) Cleavage reaction. 5’-end labelled 28-mer DNA (A) and RNA (B) treated with indicated concentrations of MsTopoI and Y339F (left and right panel respectively) and the products formed were analysed by 8 M urea-12% PAGE. (C) Protonation of the leaving group (side-chain O of Tyrosine-339) by a nearby water molecule acting as general acid (BH) would result in the release of topo I. (D) 3’-end labelled 28-mer RNA is treated with 0.05 µM of topo I in assay buffers (pH 6.0 to 9.0) and analysed using 8 M urea-12% PAGE as described in Materials and Methods. (E) Reactions with 3’-end labelled 28-mer DNA and processed as in D and Materials and Methods.

## Discussion

This work illuminates multitasking capabilities of a type IA topoisomerase that allows it to catalyse several related chemical reactions for distinct biological functions. We demonstrate that the enzyme carries out robust RNA hydrolysis in addition to topoisomerisation activities with both DNA and RNA substrates. The RNA topoisomerisation and hydrolysis activities are dependent on the active site tyrosine residue of the enzyme. We illustrate a hitherto unidentified function for topoisomerases in rRNA maturation process. The ability of the enzyme to bind and cleave tRNA and rRNA, as well as its participation in the processing of rRNA precursors, brings to light a new functional role for Type1A topoisomerases in bacteria.

The absence of topo III and topo IV in the mycobacterial genus [10, 44], combined with the large genome size (4-7 Mb), imposes extra functional burdens on the sole type IA enzyme, although some relief is provided by the stimulation of its DNA relaxing activity by factors such as HU and SSB [45, 46]. The lack of topo IV additionally calls for topo I to collaborate with DNA gyrase at the Ter region to promote decatenation and segregation of duplicated chromosome copies [17]. However, the activity of MsTopoI and MtTopoI on RNA was unsuspected, as previous studies appeared to suggest the absence of RNA topoisomerase activity for these enzymes [9]. The findings from our present study are at variance to this claim, instead conforming to the widespread presence of RNA topoisomerase activities in all three domains of life [9]. The apparent discrepancy would be due to the difference in constructs, enzyme preparation and assay conditions employed. The robust RNA binding, cleavage, strand passage and religation activities of the enzyme suggest its potential participation in multiple cellular RNA transactions. During exponential phase of growth, both rRNA and tRNA are highly transcribed. Hence, generation of RNA entanglements are not unexpected, and the topoisomerase activity of MsTopoI/MtTopoI described here may account for resolving such topological obstructions.

In search of a biological role for RNA hydrolysis activity of the enzyme, we reasoned that the *in vitro* cleavage of tRNA and rRNA by the enzyme could be non-trivial. The observation that a number of different RNases involved in processing the precursors of stable RNAs in mycobacteria is fewer than *E. coli* (Supplementary Table S1), warranted a further investigation on the *in vivo* role of topo I in rRNA processing. Under-representation of RNA processing enzymes may impose a certain burden to mycobacteria in ensuring the completion of downstream events in an orderly manner. Our results suggest that the smaller complement of such enzymes is at least partly compensated for by the topo I stepping in to assist the maturation of rRNA and tRNA. This unanticipated but remarkable RNase activity reveals a new facet of topo I catalytic repository. Although substantial cleavage is seen in the *in vitro* experiments with rRNA (and tRNA), processing should be restricted to precursor regions *in vivo*. The *in vitro* cleavage by the enzyme may represent an unregulated activity on totally exposed, protein-free RNA substrates. Such unrestricted cleavage would be attenuated *in vivo* towards processing possibly by interaction with collaborating nucleases or other regulatory proteins. A point in support of this argument is that, while topo I-mediated cleavage of both 16S and 23S rRNA is robust *in vitro*, the processing role of the enzyme *in vivo* appears to be much more prominent for 16S rRNA than 23S rRNA. This role is authenticated by the observation that topo I expression partially overcomes defective 16S rRNA processing under RNaseE depletion. The auxiliary role of topo I in RNA processing activity may be dispensable during normal growth, but it may become significant under circumstances when the processing RNase activities become inadequate. Our study thus unveils the functional collaboration between topo I and the rRNA/tRNA processing machinery in the essential steps preceding ribosome assembly and protein synthesis.

The conversion of circle to knot was used as a measure for RNA strand-passage and RNA topoisomerase activity in earlier assays [7, 9, 22]. Now, we have measured the extent of strand passage by assaying the activity and identified regions of a type IA topoisomerase participating in the step. The DNA and RNA reactions are similar in that the invariant active-site tyrosine is essential for both the RNA cleavage and religation steps, while Mg^2+^ is required for the religation step. However, as depicted in Figure 6, the RNA substrate may follow two alternative mechanistic paths, only one of which is accessible to DNA. The 2’-hydroxyl group in RNA is well-positioned to stabilize the leaving group, and RNA reactions are refractory, or partially immune, to disruption of the proton relay that blocks DNA reactions. The E21Q mutation in the His-Asp-Glu catalytic triad, which eliminates DNA cleavage by MsTopoI, permits efficient RNA cleavage and a reduced (but readily detectable) level of RNA knotting (Figure 6).

Thus, at first glance, the DNA/RNA topological reactions and RNA hydrolysis would appear to be very different activities, but from the reactions and data depicted in Figure 6 and 7, it is evident that the mechanism of RNA hydrolysis is similar to the transesterification reactions. However, unlike knotting of RNA or removal of RNA crossings, the RNA processing reaction must not only stop at the cleavage step but must also block the isoenergetic religation step; the reaction needs to be terminated with RNA hydrolysis and enzyme release. Release of the enzyme from its covalently-bound state by hydrolysis is one possible means to ensure that an RNA processing step is not aborted due to cleavage reversal. The nucleophile for the religation step is the 3’-hydroxyl group for type IA topoisomerases. This role is served by the 5’-hydroxyl group for type IB topoisomerases (and also tyrosine site-specific recombinases that follow the type IB mechanism). Alternative nucleophiles, such as water and polyhydric alcohols, are utilized by several of these enzymes for attacking the phosphotyrosyl bond of the cleaved intermediate [36, 37, 47]. During type IB topoisomerase action on an RNA substrate, the vicinal 2’-hydroxyl group can promote elimination of the enzyme from RNA linkage via the formation of a 2’, 3’-cyclic phosphodiester bond [8]. However, for type IA enzymes that form 5’ covalent complexes, the 2’-hydroxyl group is too distant from the 5’-phosphate to mediate an analogous reaction. The more likely mechanism for RNA maturation mediated by a type IA enzyme is the post-RNA cleavage hydrolysis of the 5’-phosphotyrosyl bond in a reaction akin to RNaseH activity (Figure 7). Our results (Figure 7) suggest that intracellular pH could be one factor, among others, that determines the choice between religation and hydrolysis following the formation of the RNA-topo I covalent complex.

In conclusion, this study sheds light on the surprising multi-tasking capabilities of mycobacterial topo I that include RNA tangling/untangling and RNA processing in addition to its established roles in modulating DNA topology and assisting DNA gyrase in unlinking chromosome daughter molecules for segregation [11, 16, 17, 48]. The versatility of the enzyme must have been an adaptive response to the absence of topo III and topo IV in mycobacteria in conjunction with the smaller set of RNA processing nucleases that it harbours compared to other bacteria [10, 21]. The robust RNA topoisomerase, RNA cleavage and RNA hydrolysis activities of the enzyme raise the prospect of its potential origin in an RNA world as suggested before [49]. Presumably, the ancestry of topoisomerases and site-specific recombinases goes back to an elementary RNase (or a small set of RNases), which over evolutionary time acquired the capability of DNA cleavage and religation via covalent catalysis, but did not altogether lose their competence to act on RNA. Earlier studies uncovering cryptic RNase activities in type IB topoisomerases and Flp recombinase utilized mixed RNA-DNA substrates in *in vitro* reactions [8, 50]. Our results demonstrate the hydrolysis of true RNA substrates *in vitro* and *in vivo*, validating experimentally the long-held speculation that topoisomerases could participate in RNA processing. Our findings suggest that topoisomerases could, in principle, fulfil several other biological functions that they have been speculated to participate in, for example, removal of mis-incorporated rNTPs during replication, RNA splicing and regulation of RNA folding [7, 8, 50, 51].

## Materials and Methods

### Enzymes and reagents

Wild-type and variant of MsTopoI (Y339F, E21Q, D108A and CTD mutants) were expressed and purified as described previously [16]. The glutamate (E21) residue of MsTopoI was substituted to glutamine by an inverse-PCR method [52]. After Dpn I (New England Biolabs, NEB) treatment, PCR products were transformed into *E. coli* DH5alpha cells. Positive clones were confirmed by sequencing. T4 RNA ligase, T4 polynucleotide kinase (T4 PNK), T7 RNA polymerase (T7 RNAP), RNaseH, calf intestinal phosphatase (CIP), and RNase inhibitor were from NEB. TURBO DNase I was purchased from Ambion. Terminal deoxy transferase (TDT) was obtained from ThermoFisher Scientific. Other chemicals, primers and oligonucleotides were purchased from Sigma Aldrich. Poly clonal and monoclonal antibodies against topo I were generated in the laboratory. Other reagents are laboratory reagents.

### DNA Relaxation

The assays were carried out with 500 ng of negatively supercoiled pUC18, and 0.025 to 0.1 µM of wild-type MsTopoI and variants (D108A and E21Q) in an assay buffer (buffer A, 40 mM Tris·HCl, 20 mM NaCl and 10 mM MgCl_2_) at 37°C for 30 min. The assays were also performed with 0.2 µM of MsTopoI and increasing concentrations (2.5 to 12.5 molar excess) of *E. coli* total tRNA as competitor. Reaction products were separated by electrophoresis in 1.2% agarose gels and stained with EtBr for visualization.

### Binding and cleavage

EMSAs were carried out with 5’-end labelled (with T4 PNK and γ-P^32^ ATP) initiator tRNA (itRNA, 0.01 µM), and MsTopoI and MtTopoI (0.005 to 0.04 µM) in buffer A on ice for 10 min. The products were analysed on 5% non-denaturing PAGE. For tRNA cleavage experiments, 5’-end labelled itRNA and ssDNA (10000 counts per minute (cpm), 0.01 µM) were incubated with indicated concentrations of MsTopoI and MtTopoI in buffer B (40 mM, Tris·HCl, 20 mM KCl and 1 mM EDTA). The products were analysed on 8 M urea-6% PAGE. EMSAs for rRNA were performed with MsTopoI and 500 ng of rRNA as described above and the products were separated on 1.2% agarose gel. For assessing mRNA binding, 3 and 10 µg of MsTopoI was UV cross-linked with *in vitro* transcribed body labelled mRNA (10000 cpm) and treated with 30 µg of RNaseA. The TopoI-mRNA complexes were separated on 8% SDS-PAGE followed by phosphorimaging. To visualize MsTopoI-tRNA phospho-tyrosine complexes, 3’-end labelled itRNA was generated by ligating it to 11-mer labelled ssDNA using RNA ligase. Complexes of the labelled substrate (10000 cpm, 0.01 µM) and MsTopoI (0.02 µM) were trapped with 0.5 % SDS and analysed by non-denaturing 6% PAGE. In another set of reactions, the trapped complexes were treated with 50 µg/ml proteinase K and 0.5% SDS. Some of the reactions included 50 molar excess ssDNA oligonucleotides containing a strong topoisomerase site (STS [46, 53]. To carry out cleavage assays with oligonucleotides, 28-mer DNA or RNA were 5’-end labelled withT4 PNK and ATP (γ-P^32^), and 0.01 µM of DNA/RNA (10000 cpm) was incubated with wild-type MsTopoI, and D108A and E21Q variants of MsTopoI at 37°C for 30 min in buffer B. The products were analysed by 8 M urea-12% PAGE.

To analyse RNA and DNA hydrolysis by MsTopoI, and release of the enzyme, 28-mer RNA was 3’-end labelled with pCp (5’-P^32^) and RNA ligase. The 28-mer DNA was 3’-end labelled with dATP (α-P^32^) and TDT. The labelled RNA and DNA (10000 cpm, 0.01 µM) were incubated with MsTopoI in buffers with different pH (potassium phosphate buffer pH 6.0-7.0 and Tris HCl pH 7.5 to 9.0), 1 mM EDTA, 20 mM KCl for 30 min at 37°C. The reactions were arrested with 0.5% SDS and 45% formamide, and the products were analysed by 8 M urea-12% PAGE.

### Strand passage and religation

The protocol developed for DNA strand passage [23] was adapted for RNA. MsTopoI (0.05 µM) was incubated with radiolabelled itRNA for 15 min in buffer B on ice and divided into two aliquots. To measure the initial counts, one of the aliquots was incubated with anti-topo I clamp-closing monoclonal antibody 1E4F5 [31] which locks the tRNA inside the enzyme cavity. To the second aliquot, 5 pmoles of unlabelled single stranded DNA (ssDNA) was added. The reactions were incubated for 15 min at 37°C, followed by the addition of 10 mM MgCl_2_ and further incubation for 30 min at 37°C. Next, 1E4F5 antibody was added to the second aliquot to lock the gate, followed by 30 min incubation. All the reactions were spotted on Millipore nitrocellulose filter (0.45 µm) pre-equilibrated with the buffer B, washed thrice by high salt buffer (buffer B containing 1 M NaCl), and the radioactivity retained on the filter was measured by liquid scintillation counter. Extent of strand passage was calculated as the percentage reduction in the radioactivity (cpm) **-**

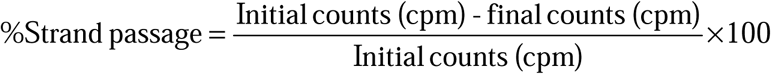

Religation assays were performed by incubating 5’-end labelled 28-mer RNA (10000 cpm, 0.01 µM) with MsTopoI (0.05 µM) at 37° C for 15 min in buffer B to obtain cleaved fragments. An unlabelled 28-mer complementary oligonucleotides (0.03 µM) was added to anneal the cleaved RNA fragments to enhance the religation, followed by the addition of MgCl_2_ (5 and 10 mM). After incubation for 15 min at 37 °C, the products were analysed by 8 M urea-12% PAGE.

### rRNA and tRNA processing

An *E. coli* KL16 strain, which is ΔRNaseR [54] was grown at 22°C for 18 hr, RNA was isolated by the Trizol (Sigma Aldrich) method [55] and integrity of the RNA was analysed on a 1.2% formaldehyde agarose gel at 100 V for 90 min. To carry out rRNA processing assay, 2 µg of total RNA was incubated with 0.2 µM of MsTopoI for 30 min in buffer B and the products were resolved on 1.2% formaldehyde agarose gel at 50 V for 150 min. Then, RNA was transferred to Nytran membrane using Biorad transfer apparatus for 150 min at 8 V, cross-linked to the membrane with 1200 KJ/cm^2^ energy of UV light. The blot was incubated overnight in prehybridization buffer containing yeast RNA and Denhardt’s solution (1% BSA, 1% Ficoll, 1% polyvinylpyrrolidone 40) at 65°C and probed for internal and precursor region of 16S and 23S rRNA [56]. 16S rRNA gene with its 3’ precursor region was PCR amplified using forward primer (17S_rRNA_F) containing T7 RNAP promoter and reverse primer (17S_rRNA_R) from *M. tuberculosis* genomic DNA. Unprocessed 16S rRNA gene was *in vitro* transcribed using T7 RNAP and was purified with phenol-chloroform extraction and ethanol precipitation. Similarly, 23S-16S intergenic region and operonic tRNA (gluT-aspT-pheU) were prepared. To carry out RNA processing assays, rRNA or tRNA transcripts (10000 cpm, 0.02 µM) were incubated with 0.005-0.160 µM of MsTopoI, MtTopoI and Y339F in buffer B and incubated at 37°C for 30 min and the products were analysed by 8 M urea - 18% and 6% PAGE, respectively.

### RNA immuno-precipitation (RIP)

*M. tuberculosis* H37Ra cells grown to OD_600 nm_ of 0.6 were treated with formaldehyde (final concentration 1%) for 10 min at 37°C. Crosslinking was quenched by the addition of glycine (final concentration 125 mM). The cell pellets were washed with 1X TBS (Tris buffered saline: 20 mM Tris·HCl pH 7.5, 200 mM NaCl), re-suspended in 4 ml RIP buffer (50 mM HEPES-KOH pH 7.5, 200 mM NaCl, 1 mM EDTA, 1% Triton X-100, 0.1% sodium deoxycholate, 0.1% SDS, protease inhibitor and RNase inhibitor, lysed in a Bioruptor (Diagenode) and debris removed by centrifugation. Mock IP and IP samples were incubated with IgG control and anti-topo I antibody respectively. Protein–RNA complexes were immuno-precipitated as described earlier [17] and RNA was extracted using the Trizol Method. cDNA was synthesized and RT-qPCR were performed to determine the enrichment of topo I on genes using specific primers (Supplementary Table S2). The experiments were carried out three times and the data represented as mean ± SEM of three independent samples.

### RNA knotting and unknotting

The 104-nucleotide single-stranded RNA was generated by *in vitro* transcription using T7 RNAP from a DNA template, adapting an approach described earlier [7]. RNA was treated with 10 units of CIP for 1 hr at 37° C, followed by phenol/chloroform extraction, ethanol precipitation and elution from the gel as described [57]. The 5’-end of the RNA was labelled with γ-P^32^ ATP and T4 PNK. A 40-mer DNA linker was used to anneal to RNA to increase the efficiency of ligation to generate circles. The ligation conditions used were: 1 unit T4 RNA ligase per µl of reaction mixture, 10% PEG-400, 50 mM Tris·HCl (pH 7.6), 10 mM MgCl_2_, 1 mM DTT, and 25 µM ATP at 16°C for 12 hr. The ligated products were treated with DNase I followed by phenol/chloroform extraction and ethanol precipitation. Formation of circles were confirmed on 8 M urea-15% PAGE. For RNA knotting, the circular RNA generated above (5000 cpm) was denatured at 90°C for 5 min in buffer C (40 mM Tris·HCl (pH 7.5), 20 mM KCl, 1 mM EDTA, 10 mM MgCl_2_ and 10 % PEG-400). MsTopoI and the active site mutant (Y339F) were added to the reaction mixtures and incubated for 90 min at 37° C. Reactions were stopped by the addition of 45% formamide followed by heating at 70°C and the products were resolved by 8 M urea-15% PAGE. To prepare entangled RNA substrate, the RNA was prepared and ligated as described above with excess of T4 RNA ligase and PEG-8000. The resulting substrate was incubated with 0.02 and 0.04 µM of MsTopoI in buffer B and products were analysed by 8 M urea-12% PAGE.

### Generation of conditional knockdown strains

An *M. smegmatis* topo I conditional knockdown strain was generated using a CRISPR interference system [19]. Guide RNA (Supplementary Table S2) was ligated to pRH2521 vector. The clones were electroporated into *M. smegmatis* MC^2^-155 containing dCas9 expression system (pRH2502). Cells were selected for hygromycin (50 µg/ml) and kanamycin (25 µg/ml) resistance. The depletion in MsTopoI level was confirmed through immuno-blotting using anti-topo I antibodies as described [14]. For the generation of an RNaseE conditional knockdown strain (RNAseE kd) of *M. smegmatis*, a 977 bp fragment from the gene encoding RNaseE was ligated to the downstream part of the Pptr promoter in a suicide vector pFRA171W [20] to obtain PptrRNAseE. To achieve the integration of the construct in the *M. smegmatis* MC^2^-155 genome, 1 μg of PptrRNAseE plasmid was electroporated and the recombinants were screened on Middlebrook 7H11 plates containing hygromycin (50 μg/ml). Integration of the construct was confirmed by PCR using vector specific primers. The depletion in RNaseE level was confirmed by RT-qPCR. For complementation with the gene of MsTopoI, the MsTopoI gene was ligated into the pMIND vector [58] and electroporated into the PptrRNaseE strain of *M. smegmatis* MC^2^-155. MsTopoI expression induced with 25 ng/ml tetracycline was assessed by immuno-blotting.

### Northern dot-blot and RT-qPCR

To analyse accumulation of precursor rRNA *in vivo* by Northern dot-blot in RNaseE knockdown strain of *M. smegmatis*, 2 µg of RNA isolated from the strain was spotted onto a Nytran membrane using a dot-blot apparatus (Biorad) and UV cross-linked (1200 energy, CL 1000 ultraviolet crosslinker). The blots were probed with 16S and 23S precursor and internal region probes (Supplementary Table S2). To examine precursor rRNA accumulation with RT-qPCR, the RNA was isolated from wild-type, MsTopoI- and RNaseE-depleted strains, and the MsTopoI complemented strain using the Trizol method [55], and the cDNA was prepared using specific reverse primers (Supplementary Table S2) for internal and precursor regions of 16S and 23S rRNA. RT-qPCR was carried out to estimate the level of precursor rRNA. The internal region was taken as control to normalize the data. The data represented is mean ± SEM of three independent experiments.

## Acknowledgements

We thank K. Drlica, A. Maxwell, J. Berger, M. Jayaram and U. Varshney for editing the MS and suggestions. We thank R. N. Husson for plasmids pRH2521 and pRH2502, R. Manganelli for plasmids pFRA171W and pFRA42A, B. Robertson for pMIND vector and U. Varshney for ΔRNaseR *E. coli* KL16 and other reagents. Yojna Dighe is acknowledged for technical assistance and the members of VN laboratory for suggestions. We acknowledge the real-time PCR and Phosphor imaging facilities of Indian Institute of Science supported by the Department of Biotechnology, Government of India.

## Conflict of interest statement

None declared.

## Notes

### Competing Interest Statement

The authors have declared no competing interest.

