## Supplemental Material for "A multifunctional type IA DNA/RNA topoisomerase with RNA hydrolysis and rRNA processing activities from *Mycobacterium smegmatis* and *Mycobacterium tuberculosis*"

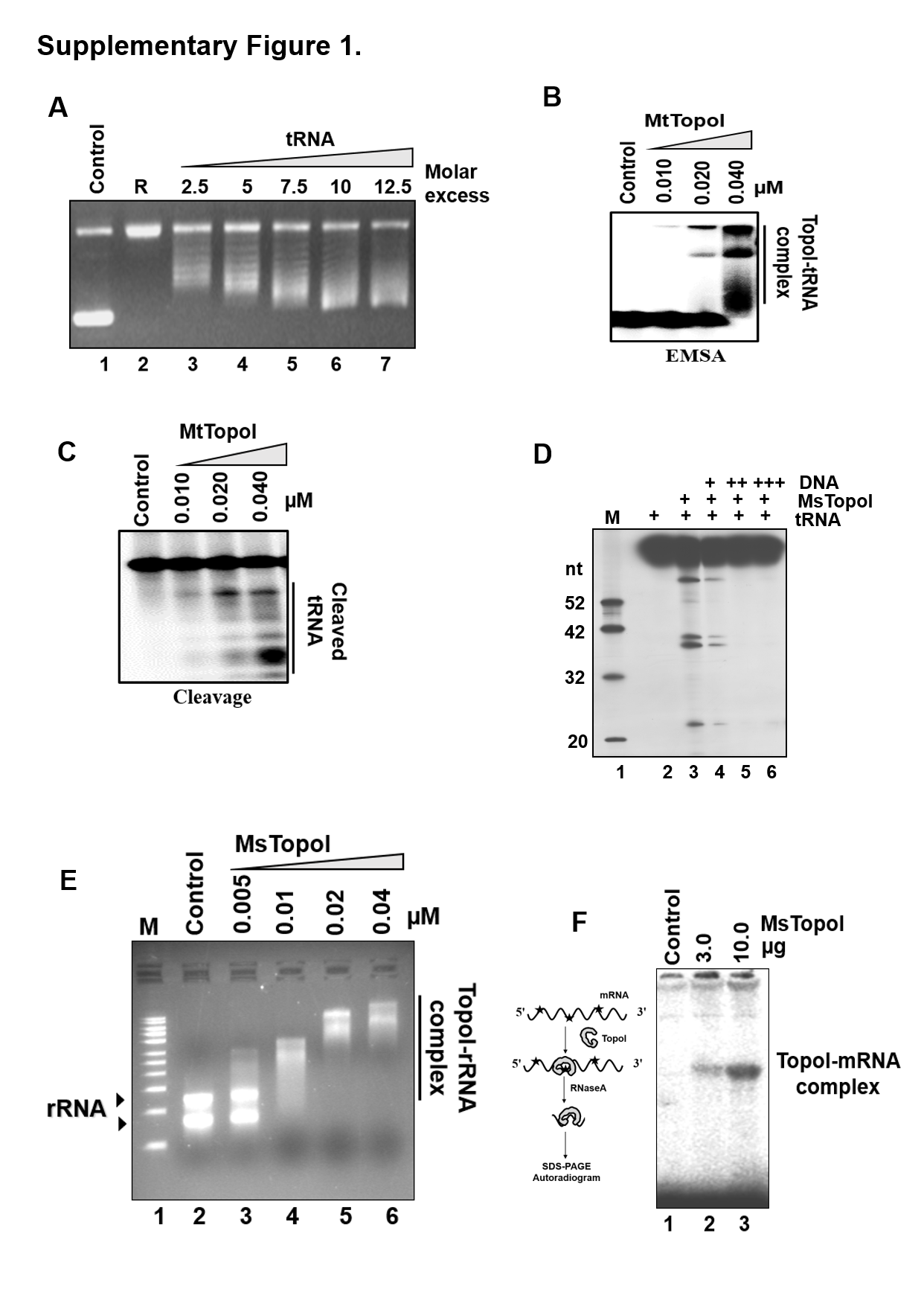

**Supplementary Figure 1.** (A) For DNA relaxation inhibition by tRNA, DNA relaxation assay was performed with MsTopoI and pUC18 in buffer A in the presence of a molar excess of *E. coli* total tRNA as described in Materials and Methods. The products were analysed on a 1.2% agarose gel and visualized after EtBr staining. Control is pUC18 without MsTopoI and R is relaxed pUC18 incubated with MsTopoI. (B and C) Binding and cleavage assays were performed with 5′-end-labelled itRNA and MtTopoI as shown. The products were analysed by 5% PAGE (B) and 8 M urea-6% PAGE (C) respectively. (D) Cleavage reactions were performed with 5´-end labelled itRNA and MsTopoI (0.05 µM) and competed by the addition of increasing concentrations of site specific (strong TopoI site, STS) ssDNA as described in Materials and Methods. (E) EMSA with rRNA. 500 ng of rRNA (purified from *E. coli* cells*)* was incubated with various concentrations of MsTopoI in buffer A (lanes 3-6). Lane 2: 500 ng of rRNA in the absence of topo I. Lane 1: 1kb ladder. (F) Interaction with mRNA. Schematic of UV crosslinking reaction is shown in the left. UV crosslinking carried out with 3 and 10 µg of topo I and *in vitro* transcribed mRNA followed by 30 µg of RNase A treatment (lanes 2-3). Lane 1: reaction in the absence of topo I. The topo I-mRNA complexes were analyzed using 8% SDS-PAGE followed by phosphorimaging.

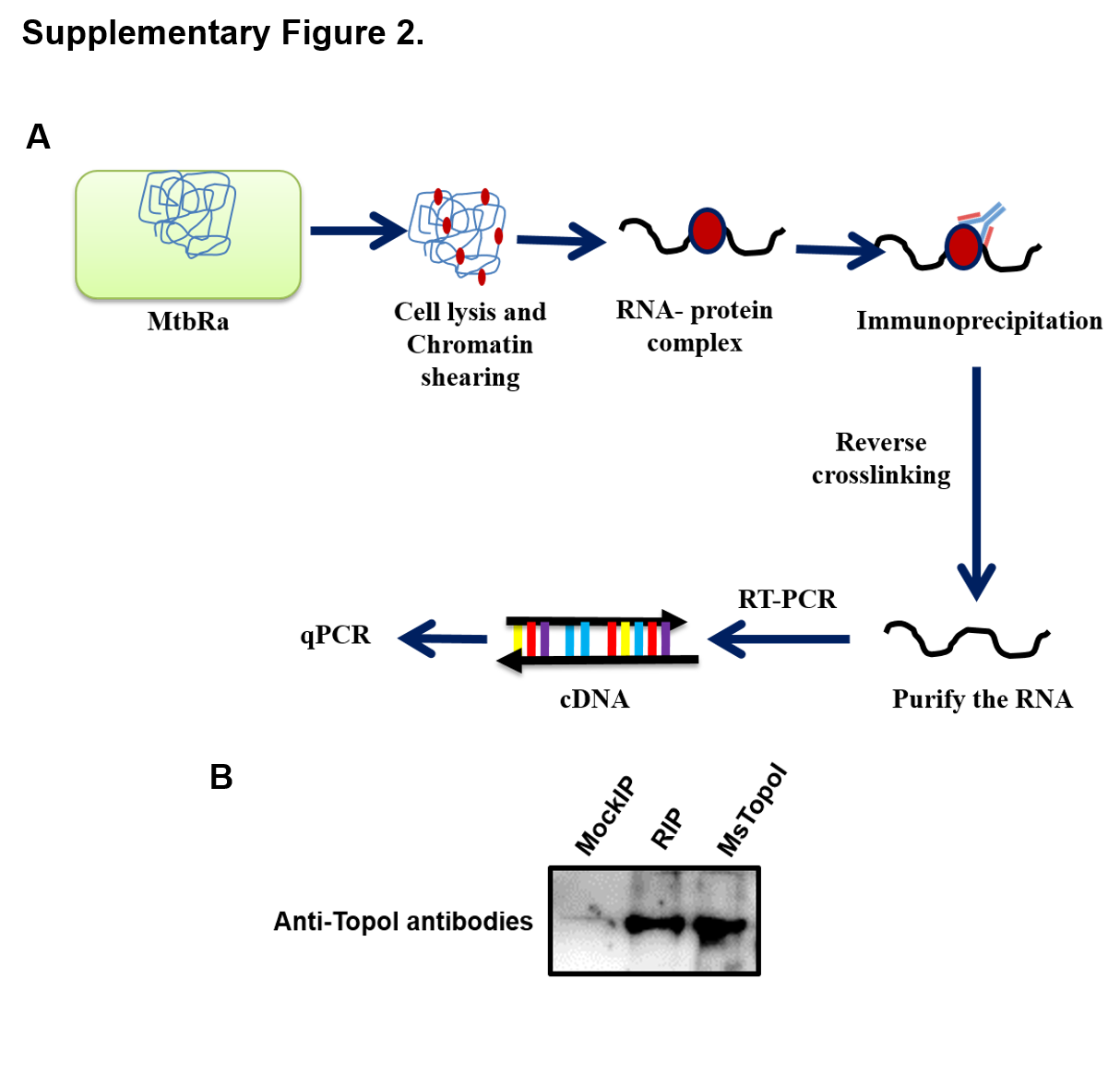

**Supplementary Figure 2.** (A) Schematic depicting the protocol for RIP-qPCR. (B) Immunoblot showing specific pull-down of TopoI in a RIP sample. Purified MsTopoI is used as marker in the last lane.

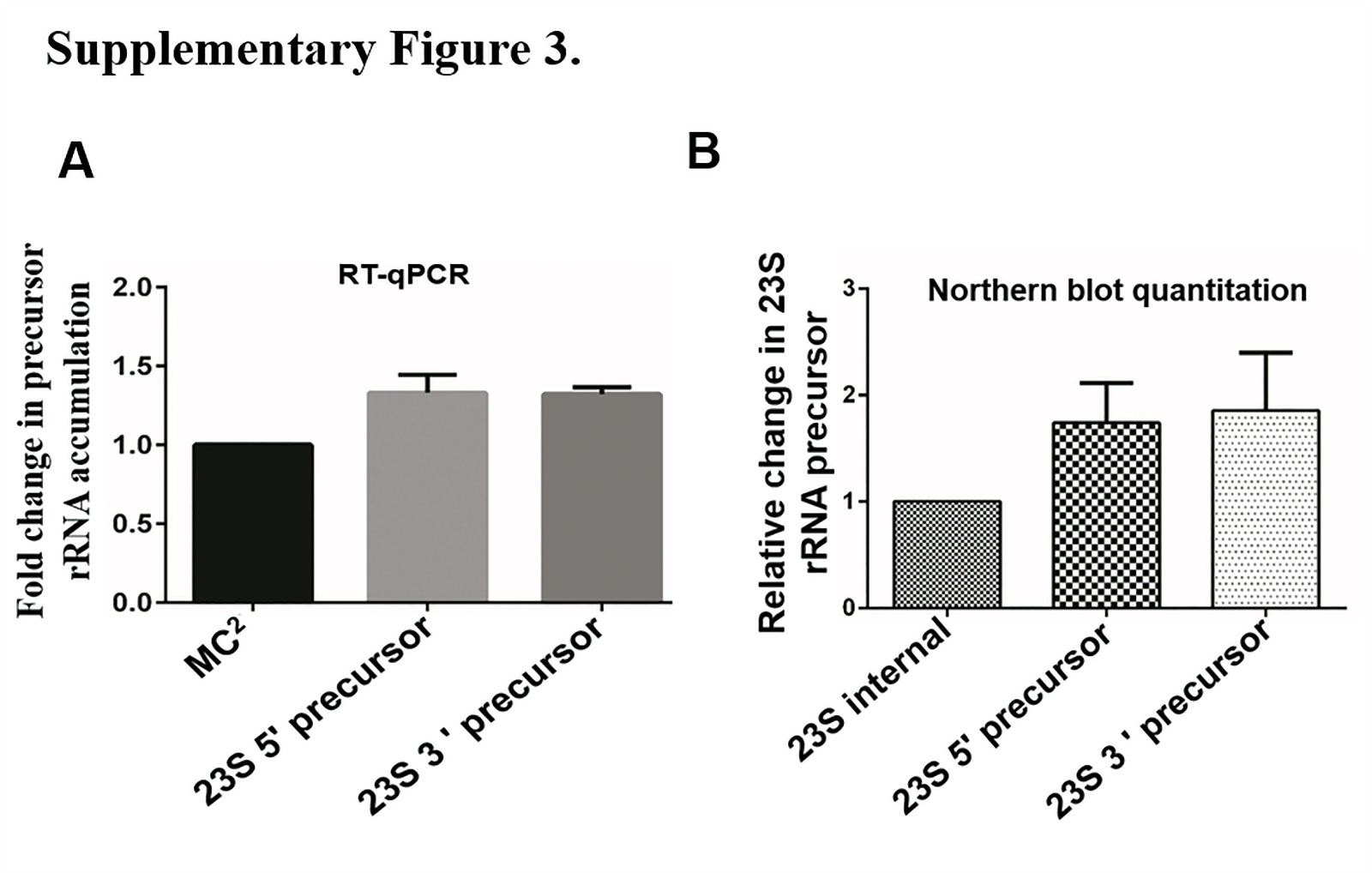

**Supplementary Figure 3.** (A) The real time quantitative PCR (RT-qPCR) performed using RT primers for 23S precursor and internal regions. The data is represented after normalizing with the internal region. Error bars represent mean ± SEM of three independent experiments. (B) Quantitation of Northern blot and data is represented in relative changes in 23S precursor rRNA. Relative fold changes were calculated by normalisation of precursor 23S rRNA band (5′ or 3′) pixel values by the internal 23S rRNA band values and expressed relative to MC^2^.

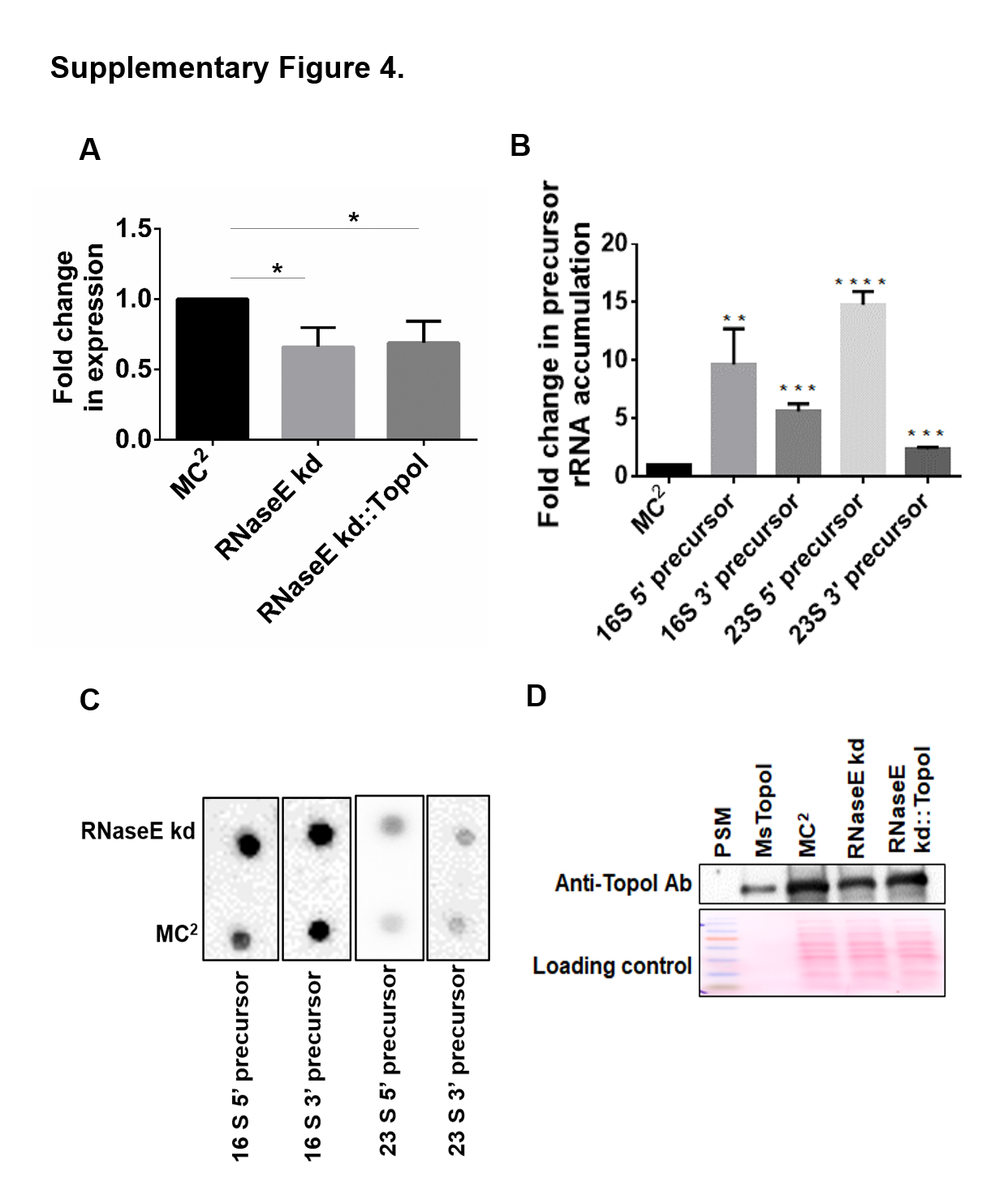

**Supplementary Figure 4.** (A) RT-qPCR to assess the RNaseE depletion in the RNaseE knockdown (RNaseE kd) strain. (B and C) Measurement of accumulation of 16S and 23S precursor rRNA in RNaseE kd strain. RNA from MC^2^ and RNaseE kd strains of *M. smegmatis* were analysed for accumulation of precursor rRNA using qPCR (B) and Northern dot-blots (C) respectively. Error bar represents Mean ± SEM values from three independent experiments. P values were determined by unpaired t test. P value≤0.05 was considered as significant. ns-non-significant, * <0.05, ** <0.01, *** <0.001 and **** <0.0001. (D) Immunoblot showing expression of MsTopoI upon complementation in RNaseE knockdown strains. Lower panel is ponceau stained blot.

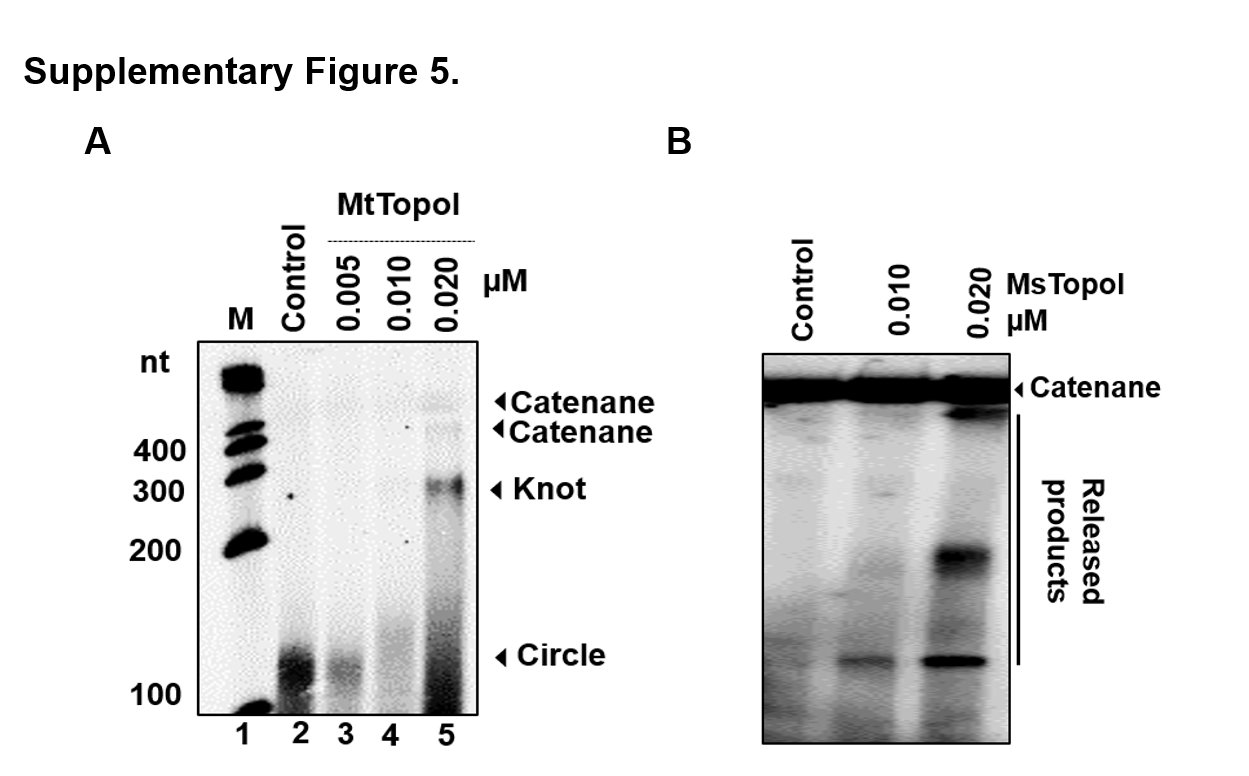

**Supplementary Figure 5.** (A) The circular RNA substrate was treated with MtTopoI and the products were analysed using 8M urea-15% PAGE. (B) Catenated RNA substrate was incubated with MsTopoI and the products were analysed using 8M urea-12% PAGE. The reaction conditions and assignment of circle/knot/catenane are according to the previous studies [1, 2].

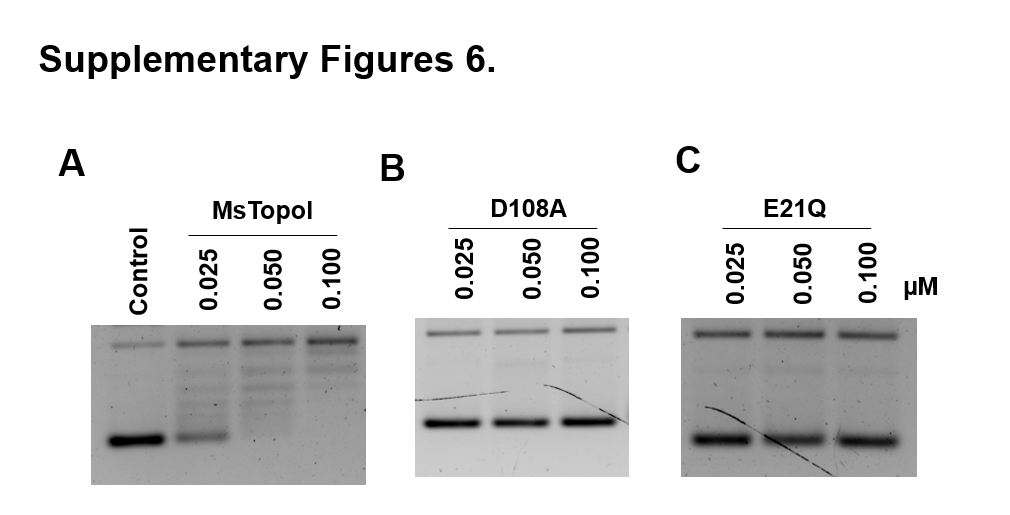

**Supplementary Figure 6.** For relaxation assays. 500 ng of pUC18 was incubated with various concentrations of MsTopoI (A), D108A (B) and E21Q (C) enzymes at 37°C for 30 min in buffer A. The products were analyzed on 1.2% agarose gel as described in Materials and Methods. The gels were stained with EtBr.

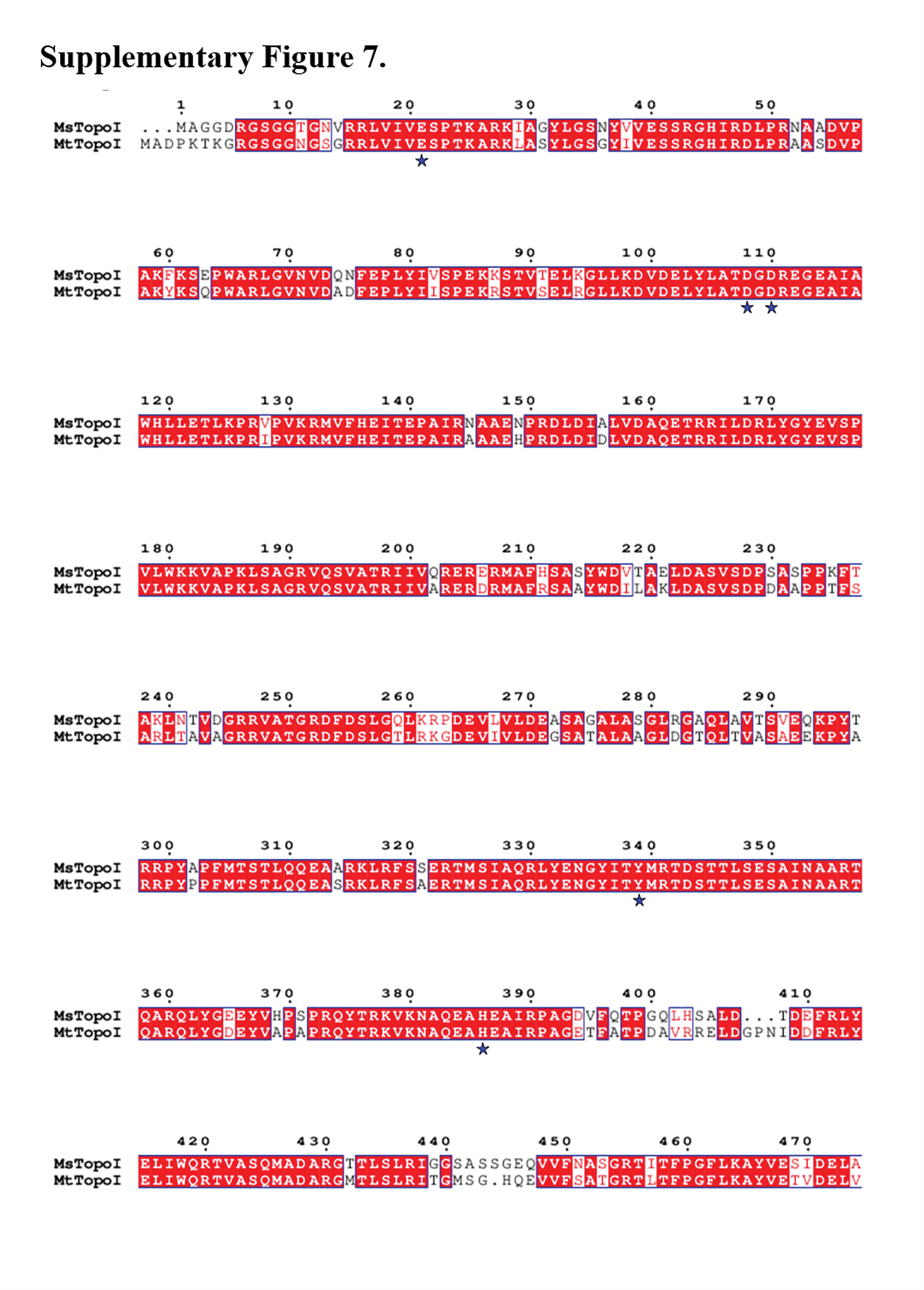

**Supplementary Figure 7.** The sequence alignment corresponding to N-terminus 500 residues of topo I from *M. tuberculosis* and *M. smegmatis* is shown. Common residues implicated for the topo I mechanism are indicated by blue stars on the alignment. Conserved residues are marked in white on a red background while similar residues are in red. Residues in blue frame depict similarity across groups. Residue 339 is the active site tyrosine in MsTopoI and corresponds to 342 of MtTopoI.

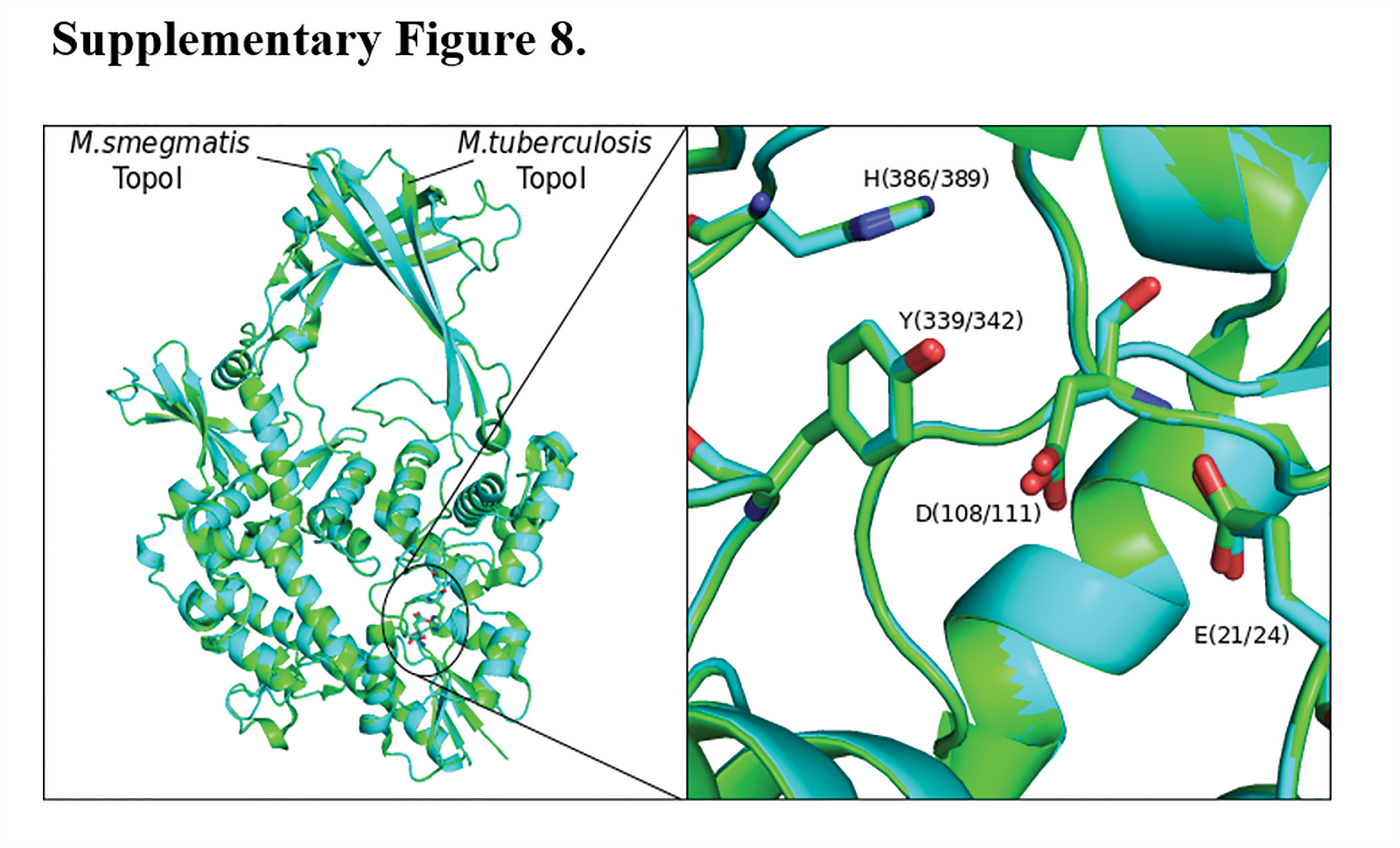

**Supplementary Figure 8.** Superposition of homology modelled structure of *M. smegmatis* TopoI (cyan) with the crystal structure (PDB ID 5D5H) of *M. tuberculosis* TopoI (green) [3] and close-up view of residues implicated for topo I mechanism. Amino acid residues involved in catalysis are indicated (*M. smegmatis/M. tuberculosis)*.

**Supplementary Table S1: Ribonucleases required for maturation of RNA from *E. coli* and mycobacteria**

| **S.No.** | **Ribonuclease (endo/exo)** | ***E. coli*** | ***M. smegmatis*** | ***M. tuberculosis*** |
| --- | --- | --- | --- | --- |
| **1.** | **RNaseE (endo)** | **+** | **+** | **+** |
| **2.** | **RNaseG (endo)** | **+** | **-** | **-** |
| **3.** | **RNaseIII (endo)** | **+** | **+** | **+** |
| **4.** | **RNaseP (endo)** | **+** | **+** | **+** |
| **5.** | **RNaseZ (endo)** | **+** | **+** | **+** |
| **6.** | **RNaseI (endo)** | **+** | **-** | **-** |
| **7.** | **RNaseJ (endo/exo)** | **-** | **+** | **+** |
| **8.** | **PNPase (exo)** | **+** | **+** | **+** |
| **9.** | **RNaseT (exo)** | **+** | **-** | **-** |
| **10.** | **RNaseR (exo)** | **+** | **-** | **-** |
| **11.** | **RNaseII (exo)** | **+** | **-** | **-** |
| **12.** | **Orn (exo)** | **+** | **+** | **+** |

**Supplementary Table S2: Oligonucleotides used for the study**

| Sr. No. | Oligonucleotide name | Sequence (5´-3´) |
| --- | --- | --- |
| 1 | rrsA_upstream_F | TCACCTGTCTTTGGATGGGT |
| 2 | rrsA_upstream_R | TGCATGTGTTAAGCACGCC |
| 3 | rrsA_downstream_F | CCGTCACGTCATGAAAGTCG |
| 4 | rrsA_downstream_R | TGTCTCTCGTGGTGCTCCTT |
| 5 | rrsA_internal_F | CCACACTGGGACTGAGATAC |
| 6 | rrsA_internal_R | TTCTTCTGCACATACCGTCA |
| 7 | rrsA_5’ precursor_Northern | CAACCCATCCAAAGACAGGTGAATTC |
| 8 | AtpB_FP | GTCGGACACCCGATCAAGTT |
| 9 | Atp_RP | GAAAAGTCGGAGCGACAACG |
| 10 | 23S_upstream_F | GTTTTTTGTGTTGTAAGTGTTTAAG |
| 11 | 23S_upstream_R | TGACACGACATCACTCGTG |
| 12 | 23S_downstream_F | GATAAGGCCCCCCGCAGACCACG |
| 13 | 23S_downstream_R | GGTTGCGAGCGCGGTGCGTCAATG |
| 14 | 23S_internal_F | CGTGCGCTTACAATCCGTCAGA |
| 15 | 23S_internal_R | TGTGTGGATACGCCCTATTCAG |
| 16 | 23S_5’ precursor_Northern | CTTAAACACTTACAACACAAAAAACC |
| 17 | (NsiI) RNaseE_pptr_FP | CCAATGCATTCCAGCCATCATTCAGCCAG |
| 18 | (ClaI) RNaseE_pptr_RP | CCATCGATGGAGTCCTCACCTGACTCCT |
| 19 | tRNA operon_F | ACTTCGAAATTAATACGACTCACTATAGCCCCCTTCGTCTAGACGGC |
| 20 | tRNA operon_R | TGGCCAGGGGCGGGATCGAAC |
| 21 | MsTopoI_E21-Q_Fp | CGACTCGTGATTGTCCAGTCGCCGACGAAG |
| 22 | MsTopoI_E21-Q_Rp | CTTCGTCGGCGACTGGACAATCACGAGTCG |
| 23 | Ms_TopA_sg1_FP | GGGATGCGGGGCAGGTCACGGATG |
| 24 | Ms_TopA_sg1_RP | AAACCATCCGTGACCTGCCCCGCA |
| 25 | Without STS | ACTTCGAAATTAATACGACTCACTATAGAGGTTTTTTTTCAGGTCCAGTCTTTTTTTTTTTTGTCAGACGGATCTTTTTTTTTTTTGACTGGACCTGATTTTTTTTTTTTGATCCGTCTGACTTTTTTGAT |
| 26 | Without STSc | TATTTCGAAGATTATGCTGAGTGATATCTCCGAGAAAAAGTCCAGGTCAGAAAAAAAAAAAACAGTCTGCCTAGAAAAAAAAAAAACTGACCTGGACTAAAAAAAAAAAACTAGGCAGACTGAAAAAACTA |
| 27 | Without STS linker | CTAGGCAGACTGAAAAAACCCGAGAAAAAGTCCAGGTCAG |
| 28 | FabD_FP | GGTTCGCAAACCGAGGGAAT |
| 29 | FabD_RP | TCTAGATCAGCGGCTTTCGAC |
| 30 | 17S_rRNA_F | ACTTCGAAATTAATACGACTCACTATAGGTTTTGTTTGGAGAGTTTGATC |
| 31 | 17S_rRNA_R | CTTACAAAACACAACAAAAACCAAAG |
| 32 | 16S-23S rRNA int_F | ACTTCGAAATTAATACGACTCACTATAGAAGGAGCACCACGAAAACGC |
| 33 | 16S-23S rRNA int_R | CTTACAAAACACAACAAAAACCAAAG |
| 34 | gyrA_FP | TCGATCTACGACAGCCTGG |
| 35 | gyrA_RP | GCTTCGGTGTACCTCATCG |
| 36 | gyrB_FP | GGACGCACCAGGAAGAAAG |
| 37 | gyrB_RP | GGTCGCACACCACTGTACC |
| 38 | gre_FP | AGGTGTACTACAACGGCGAC |
| 39 | gre_RP | CGTGTAGCTGCGGGTCTC |
| 40 | 16SrRNA_FP | ACGCGAAGAACCTTACCTGG |
| 41 | 16SrRNA_RP | CGTGCGCTTACAATCCGTCAGA |
| 42 | Pheu-tRNA_FP | GGTACCCCCTACGGGATTCG |
| 43 | Pheu-tRNA_RP | GCCCCCTTCGTCTAGACGGC |
| 44 | 16S 5' precursor | CGACGTTAAGAATCCGTATCTTCGAGTGCC |
| 45 | 16S 3' precursor | TGTGTGAGCACTGCAAAGAACGCTTTAAGG |
| 46 | 16S internal | TCTTCGCGTTGCATCGAATT |
| 47 | 23S 5' precursor | CGCTTAACCTCACAAC |
| 48 | 23S 3' precursor | GGCGTTGTAAGGTTAA |
| 49 | 23S internal | CGCGCAGGCCGACTCGACCAGTGAGC |
